# Ether Lipid Biosynthesis Promotes Lifespan Extension and Enables Diverse Prolongevity Paradigms in *Caenorhabditis elegans*

**DOI:** 10.1101/2021.09.02.457410

**Authors:** Lucydalila Cedillo, Sainan Li, Fasih M. Ahsan, Nicole L. Stuhr, Yifei Zhou, Yuyao Zhang, Adebanjo Adedoja, Luke M. Murphy, Armen Yerevanian, Sinclair Emans, Khoi Dao, Zhaozhi Li, Nicholas D. Peterson, Jeramie Watrous, Mohit Jain, Sudeshna Das, Read Pukkila-Worley, Sean P. Curran, Alexander A. Soukas

## Abstract

Biguanides, including the world’s most commonly prescribed drug for type 2 diabetes, metformin, not only lower blood sugar, but also promote longevity in preclinical models. Epidemiologic studies in humans parallel these findings, indicating favorable effects of metformin on longevity and on reducing the incidence and morbidity associated with aging-related diseases. In spite of this promise, the full spectrum of molecular effectors responsible for these health benefits remains elusive. Through unbiased screening in *C. elegans*, we uncovered a role for genes necessary for ether lipid biosynthesis in the favorable effects of biguanides. We demonstrate that biguanides prompt lifespan extension by stimulating ether lipid biogenesis. Loss of the ether lipid biosynthetic machinery also mitigates lifespan extension attributable to dietary restriction, target of rapamycin (TOR) inhibition, and mitochondrial electron transport chain inhibition. A possible mechanistic explanation for this finding is that ether lipids are required for activation of longevity-promoting, metabolic stress defenses downstream of the conserved transcription factor Nrf2/*skn-1*. In alignment with these findings, overexpression of a single, key, ether lipid biosynthetic enzyme, *fard-1*/FAR1, is sufficient to promote lifespan extension. These findings illuminate the ether lipid biosynthetic machinery as a novel therapeutic target to promote healthy aging.

## Introduction

Metformin is first line therapy for type 2 diabetes and the most frequently prescribed oral hypoglycemic medication worldwide (Inzucchi et al., 2012). Human epidemiologic studies note an association between metformin use and decreased incidence of cancer (Evans et al., 2005; Yuan et al., 2013). In addition, metformin extends lifespan in invertebrate and vertebrate models (Cabreiro et al., 2013; Martin-Montalvo et al., 2013; Onken and Driscoll, 2010), and therefore may reduce aging-related diseases in humans (Barzilai et al., 2016). Nonetheless, our understanding of the molecular pathways governing the health-promoting effects of metformin is only just beginning to emerge. Our previous work identified a conserved signaling axis connecting mitochondria, the nuclear pore complex, and mTORC1 inhibition that is required for metformin-mediated extension of lifespan in *C. elegans* and inhibition of growth in worms and human cancer cells (Wu et al., 2016). The energy sensor AMP-activated protein kinase (AMPK) is not necessary for metformin-induced growth inhibition in *C. elegans* but is required for the drug’s prolongevity effects (Cabreiro et al., 2013; Chen et al., 2017; Onken and Driscoll, 2010). Consistently, mechanistic studies indicate that the longevity-promoting transcription factor Nrf2/*skn-1* is required for biguanide-mediated lifespan extension (Cabreiro et al., 2013; Onken and Driscoll, 2010). The relationship of these metformin longevity response elements to each other and their hierarchy in the biguanide response other remains unknown. Thus, the mechanisms by which metformin exacts its beneficial effects on health are likely to be branching and complex.

The importance of ether lipids, a major structural component of cell membranes, to aging and longevity is not fully established. Ether lipids are involved in the maintenance of general membrane fluidity and in the formation of lipid rafts within microdomains, which are important for promotion of membrane fusion and cellular signaling (Glaser and Gross, 1994; Komljenovic et al., 2009; Marrink and Mark, 2004). Ether lipids have broad roles in the regulation of cell differentiation (Davies et al., 2001; Facciotti et al., 2012; Rodemer et al., 2003; Teigler et al., 2009), cellular signaling (Albert et al., 2003; Thukkani et al., 2002), and reduction of oxidative stress through their action as antioxidants (Maeba et al., 2002; Morand et al., 1988; Reiss et al., 1997; Zoeller et al., 1988). Humans deficient in ether lipid biogenesis suffer from rhizomelic chondrodysplasia punctata (RCDP), a rare genetic disorder, which results in skeletal and facial abnormalities, psychomotor retardation, and is uniformly fatal typically before patients reach their teenage years (White et al., 2003). Thus, evidence linking alterations in ether lipid levels to aging and longevity in humans is strictly correlative (Gonzalez-Covarrubias et al., 2013; Pradas et al., 2019).

Ether lipids, which are structurally distinct from canonical phospholipids, have a unique biosynthetic pathway through which a fatty alcohol is conjugated to the glycerol backbone at the *sn-1* position via an ether linkage. Ether lipid precursors are first synthesized by enzymes associated with the membranes of peroxisomes (Ghosh and Hajra, 1986; Hardeman and Van den Bosch, 1989; Singh et al., 1993). The main enzymes involved in ether lipid biosynthesis within the peroxisomal matrix are glyceronephosphate O-acyltransferase (GNPAT) and alkylglycerone phosphate synthase (AGPS). Fatty acyl-CoA reductase 1 (FAR1) supplies most of the fatty alcohols used to generate the ether linkage in the precursor, 1-O-alkyl-glycerol-3-phosphate. This precursor is then trafficked to the endoplasmic reticulum (ER) for acyl chain remodeling to produce various ether lipid products (Hua et al., 2017). In *C. elegans*, loss-of-function mutations of any of the three main enzymes involved in human ether lipid biosynthesis, *acl-7*/GNPAT, *ads-1*/AGPS, and *fard-1*/FAR1, results in an inability to produce ether linked lipids, as in humans, and has been reported to shorten lifespan (Drechsler et al., 2016; Shi et al., 2016). Worms and human cells deficient in ether lipids exhibit compensatory changes in phospholipid species, including increases in phosphatidylethanolamines and phosphatidylcholines containing saturated fatty acids (Benjamin et al., 2013; Rodemer et al., 2003). However, in contrast to humans, ether lipid deficient nematodes develop to adulthood at a normal rate, providing an opportunity to determine the biological roles of ether lipids in aging and longevity without pleiotropies associated with developmental rate.

Here, we show that the ether lipid biosynthetic machinery is necessary for lifespan extension stimulated by metformin or the related biguanide phenformin in *C. elegans*. Metabolomic analysis indicates that phenformin treatment drives increases in multiple phosphatidylethanolamine-containing ether lipids. Interestingly, requirement for the ether lipid biosynthetic genes extends to multiple genetic longevity paradigms including defective mitochondrial electron transport function (*isp-1*), defective pharyngeal pumping/ caloric restriction (*eat-2*), and compromises in mTOR complex 1 activation (*raga-1*). We show that overexpressing *fard-1*, the enzyme that supplies all the fatty alcohols for ether lipid biogenesis in *C. elegans*, extends lifespan, supportive of the idea that alterations in the ether lipid landscape alone is sufficient to promote healthy aging. Mechanistically, ether lipids promote longevity downstream of biguanide action through activation of metabolic stress defenses through the transcription factor *skn-1*/*Nrf2*. These data suggest that a heretofore unappreciated role for ether lipids is to promote organismal-level, longevity-promoting stress defenses.

## Results

### Genes responsible for ether lipid biosynthesis are necessary for biguanide-induced lifespan extension

A screen of ∼1000 metabolic genes for RNA interference (RNAi) knockdowns that interfere with the growth-inhibitory properties of metformin in *C. elegans* (Wu et al., 2016), yielded *fard-1* and *acl-7*, which are required for ether lipid biosynthesis. Ether lipids are distinguished from canonical phospholipids as the latter contain exclusively fatty acids conjugated to glycerol, whereas ether lipids contain a fatty alcohol conjugated to the glycerol backbone at the *sn-1* position via an ether linkage (Figure 1A). Confirming our screen results, quantitative analysis following RNAi knockdown of *fard-1* and *acl-7* results in significant resistance to biguanide-induced growth inhibition (Figure 1—figure supplement 1A). Our lab has previously demonstrated that biguanide effects on growth in *C. elegans* share significant overlap mechanistically with the machinery by which metformin extends lifespan in the worm (Wu et al., 2016). Indeed, loss-of-function mutations in any of three genes encoding enzymes required for ether lipid biosynthesis, *fard-1, acl-7* or *ads-1*, completely abrogates lifespan extension induced by both metformin and the related biguanide phenformin (Figure 1B—G, and throughout manuscript see Supplementary file 1 for all tabular survival statistics and biological replicates). Confirming that these mutations confer resistance to metformin by compromising ether lipid synthetic capacity, RNAi knockdowns of *fard-1* and *acl-7* also partially impair lifespan extension promoted by phenformin (Figure 1—figure supplement 1B—C). Studies from this point forward are presented predominantly with phenformin because phenformin is more readily absorbed without need for a specific transporter, unlike metformin (Segal et al., 2011; Sogame et al., 2009; Wu et al., 2016), and our experience indicates more consistent lifespan extension with phenformin in *C. elegans*.

**Figure 1.**
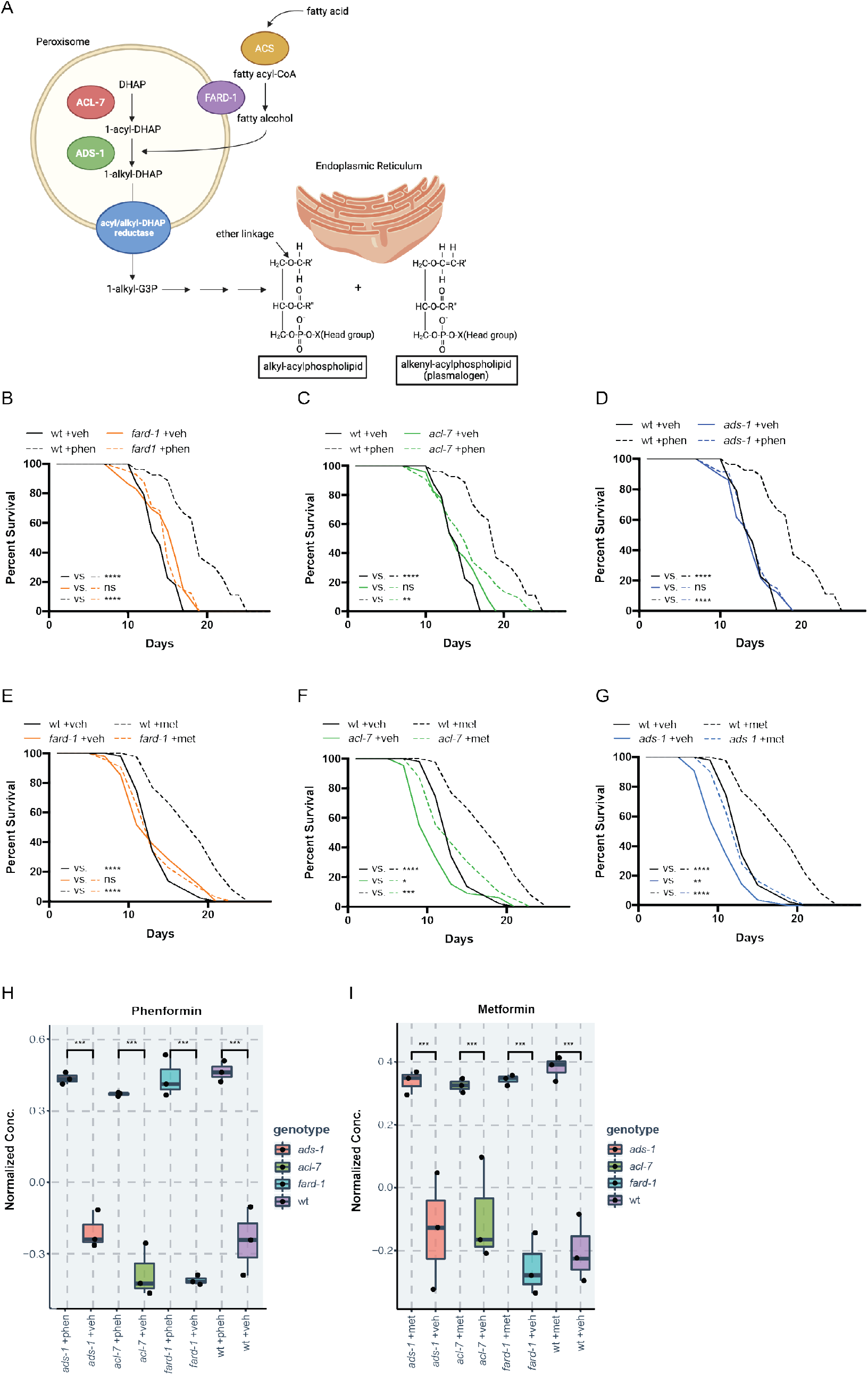
Genes responsible for ether lipid biosynthesis are necessary for biguanide-induced lifespan extension. (A) *C. elegans* ether lipid synthesis is catalyzed by three enzymes: fatty acyl reductase FARD-1, acyltransferase ACL-7 and alkylglycerone phosphate synthase ADS-1 (adapted from (Dean and Lodhi, 2018; Shi et al., 2016)). The latter two are localized to the peroxisomal lumen. (B-D) Missense, loss of function mutations in *fard-1* (B), acl-7 (C), and *ads-1* (D) in *C. elegans* suppress phenformin-induced lifespan extension. (E-G) A deficiency of ether lipid synthesis in *fard-1* (E), *acl-7* (F), and *ads-1* (G) worm mutants blunts metformin-induced lifespan extension. Results are representative of 3 biological replicates. *, *P* < 0.05; **, *P* < 0.01; ***, *P* < 0.001; ****, *P* < 0.0001 by log-rank analysis. See also Figure 1-figure supplement 1 and refer to Supplementary file 1 for tabular survival data and biological replicates. (H-I) Normalized concentrations of phenformin (H) and metformin (I) in vehicle, 4.5mM phenformin, or 50 mM metformin treated wild type (wt) *C. elegans* versus *fard-1, acl-7*, and *ads-1* mutants. n = 3 biological replicates; ***, *P* < 0.004 by two-tailed students *t*-test with Bonferroni correction for multiple hypothesis testing. Box represents 75^th^/25^th^ percentiles, while whisker represents higher/lower hinge +/-[1.5 * interquartile range (IQR)].

Because ether lipids are a major structural component of cell membranes, one possibility is that deficiencies in ether lipid synthesis compromises drug action by reducing biguanide bioavailability in the worm. To test this, we compared the relative levels of biguanides present in vehicle- and biguanide-treated wild type to the three ether lipid synthesis mutants by liquid chromatography—tandem mass spectrometry (LC-MS/MS). A comparison of normalized concentrations of phenformin across all four strains shows that phenformin abundance is quantitatively similar across wild type and the three, ether lipid mutant strains (Figure 1H and Figure 1—figure supplement 1D). Similar results were obtained when comparing levels of metformin wild type and ether lipid mutant animals (Figure 1I and Figure 1—figure supplement 1E). Thus, a deficiency in ether lipid synthesis does not significantly impact levels of biguanides in *C. elegans*.

### Phenformin induces changes in ether lipid levels

We reasoned that if biguanides require ether lipid biosynthesis to promote lifespan extension that phenformin may promote synthesis of one or more ether lipids. To investigate the impact of biguanides on ether lipids at a high level, we first utilized gas chromatography-mass spectrometry (GC-MS) analysis. We first recapitulated the observation that *fard-1* mutants show absence of 18-carbon containing fatty acid derivatives (dimethylacetals, or DMAs, which indicate alkenyl ether lipid or plasmalogen levels) and an accumulation of stearate (18:0) relative to wild type controls by GC-MS (Figure 2A—B) (Shi et al., 2016). We then asked if phenformin impacts the levels of 18-carbon alkenyl ether lipids in wild type animals and if those corresponding changes are absent in *fard-1* mutants. Strikingly, phenformin-treated wild type worms display a significant increase in 18:0 DMA versus vehicle, whereas no such increase is evident in drug treated *fard-1* worms (Figure 2C). In addition, relative proportions of stearic acid (18:0) levels within the total fatty acid pool are significantly increased in *fard-1* mutants treated with phenformin versus vehicle treated *fard-1* controls (Figure 2D). In comparison, the relative proportion of stearic acid does not rise in phenformin treated wild type animals, suggesting that stearate is being utilized for ether lipid production. Analysis of the total fatty acid pool by GC-MS (Figure 2—figure supplement 1) indicates that aside from several fatty acids (e.g. 18:2), the most pronounced differences were in the plasmalogen pool. In alignment, an assessment of levels of additional alkenyl fatty alcohols in phenformin-treated, wild type animals indicates a parallel, significant increase in the less abundant 16:0 DMA and 18:1 DMA species (Figure 2E). We conclude that phenformin treatment leads to an overall increase of alkenyl ether lipid levels in *C. elegans*.

**Figure 2.**
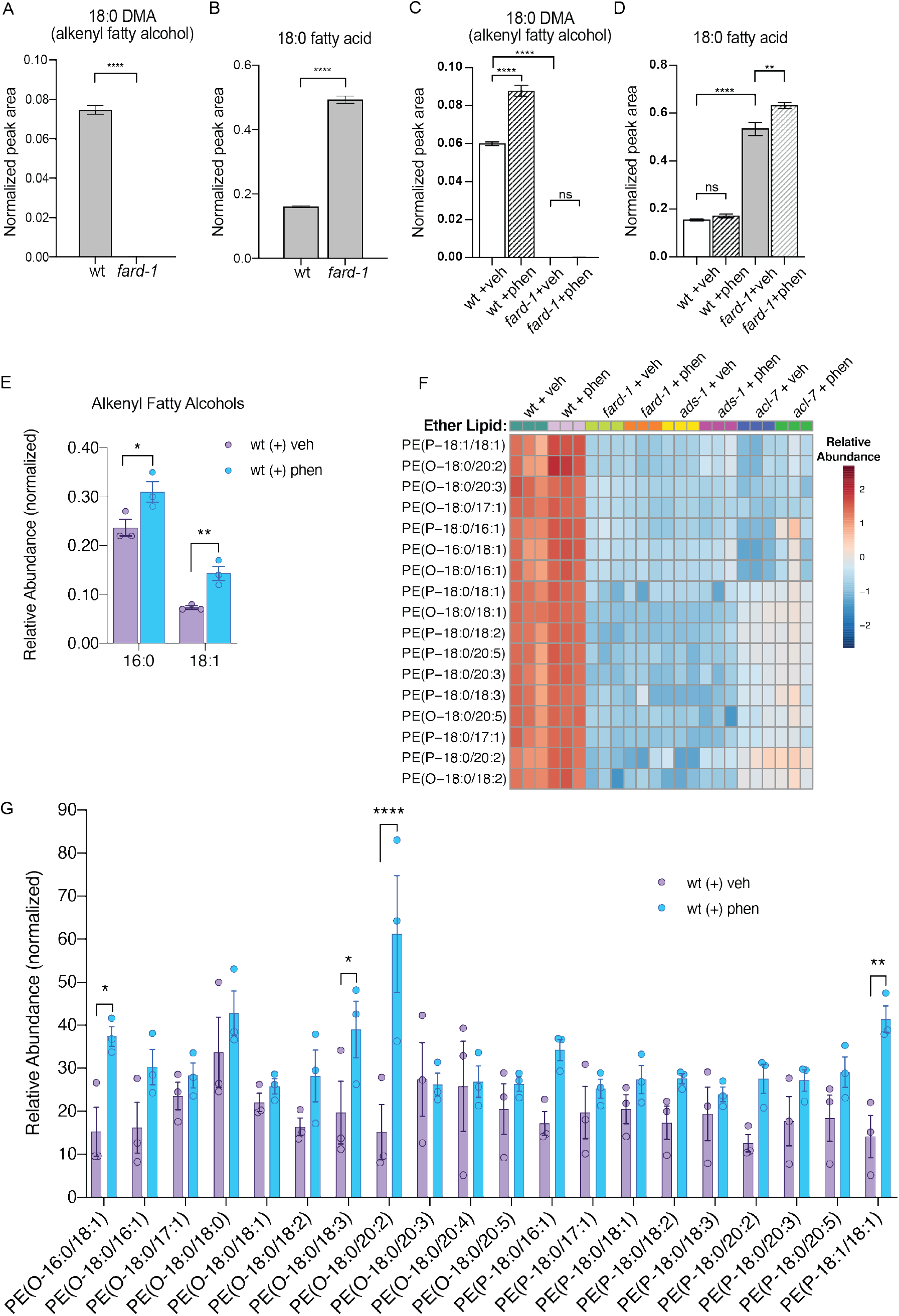
Phenformin treatment of *C. elegans* leads to increased abundance of multiple alkyl and alkenyl ether lipids. (A-B) Loss-of-function *fard-1* mutants have significant reduction in 18:0 fatty alcohols derivatized from 18-carbon containing alkenyl ether lipids (dimethylacetal, DMA) by GC/MS (A) and accumulation of the saturated fatty acid stearate (18:0, B). (C) Wild type worms treated with 4.5 mM phenformin display a significant increase in 18:0 DMA relative to vehicle control, indicative of higher levels of alkenyl ether lipids, with levels remaining essentially undetectable in *fard-1* mutants on vehicle or drug. (D) 4.5mM phenformin treatment does not impact stearate levels in wild type worms, however it does result in a greater accumulation of stearate in *fard-1* mutants. For A-D, **, *P* < 0.01; ****, *P* < 0.0001, by t-test (A-B) or two-way ANOVA (C-D), n = 3 biological replicates. (E) 4.5mM phenformin treatment results in a significant increase in 16:0 DMA and 18:1 DMA in wild type worms, relative to vehicle-treated controls (*, *P* < 0.05; **, *P* < 0.01, by multiple t-tests, with two-stage linear step-up procedure of Benjamini, Krieger and Yekutieli. n = 3 biological replicates. (F) Heat map of normalized ether lipid abundance following phenformin treatment in wild type *C. elegans* indicates an overall increase in ether lipids relative to vehicle treated controls, and this shift is absent in ether lipid deficient mutants. All metabolites shown have an FDR adjusted P < 0.05 by one-way ANOVA followed by Fisher’s LSD post-hoc testing for WT versus *fard-1, ads-1*, and *acl-7* mutants. (G) LC-MS analysis shows that phosphatidylethanolamine-containing ether lipids detected exhibited a general trend towards increased abundance in wild type worms treated with 4.5mM phenformin. Four of these ether lipids reached statistical significance: PE(O-16:0/18:1), PE(O-18:0/18:3), PE(O-18:0/20:2), and PE(P-18:1/18:1). Eleven of the ether lipids detected are of the alkyl-type (indicated by “O” in their name prior to fatty alcohol designation) whereas 9 are of the alkenyl-type (plasmalogen, indicated by “P” in their name prior to the fatty alcohol designation) ether lipids. For *G*, *, *P* < 0.05; **, *P* < 0.01; ****, *P* < 0.0001, by multiple t-tests, with multiple hypothesis testing correction by two-stage step-up method of Benjamini, Krieger, and Yekutieli, n = 3 biological replicates. See Supplementary file 2 for raw and normalized mass spectrometry data.

To investigate relative changes in individual ether lipid abundance in response to phenformin at high resolution, we utilized LC-MS analysis. Using this method, we detected 20 alkyl and alkenyl phosphatidylethanolamine-based ether lipids previously noted to be the most abundant ether lipids in *C. elegans* (Drechsler et al., 2016; Shi et al., 2016) (Figure 2F—G and Supplementary file 2). This analysis indicates that phenformin treatment results in a significant increase in normalized abundance of four ether lipids, PE(O-16:0/18:1), PE(O-18:0/18:3), PE(O-18:0/20:2), and PE(P-18:1/18:1), even when corrected for multiple hypothesis testing. The vast majority of ether lipids measured display mean levels that increase with phenformin treatment, though most are either nominally significant or exhibit a nonsignificant trend because of the strict threshold required to reach significance when correcting for multiple hypotheses. Finally, phosphatidylethanolamine ether lipid abundances were extremely low in *fard-1, acl-7* and *ads-1* mutants and unchanged by phenformin treatment, unlike in wild type animals (Figure 2F and Supplementary file 2). In aggregate, these data indicate that phenformin treatment leads to increased abundance of multiple ether lipid species in *C. elegans*.

### Peroxisomal ether lipid synthesis is essential to phenformin effects

In order to begin to understand the governance of ether lipid biosynthesis by biguanides, we examined expression of a *C. elegans* FARD-1::RFP translational reporter, under the control of its own promoter (Figure 2—figure supplement 2A). Exogenously expressed FARD-1 *(fard-1 oe1)* is expressed in intestine and localizes near structures resembling lipid droplets by Nomarski microscopy (Figure 2—figure supplement 2B). Given that ether lipid biogenesis occurs between peroxisomes and the endoplasmic reticulum (ER) (Ghosh and Hajra, 1986; Hardeman and Van den Bosch, 1989; Hua et al., 2017; Singh et al., 1993), we crossed this FARD-1::RFP reporter to an animal bearing a GFP reporter that illuminates peroxisomes in intestine (GFP fused to a C-terminal peroxisomal targeting sequence 1 (PTS1)) to determine if localization of FARD-1 is regulated by biguanides. FARD-1 does not possess a predicted C-terminal PTS1 as do ACL-7 and ADS-1. At baseline, FARD-1::RFP fluorescence partially overlaps with peroxisomally-targeted GFP (Figure 2—figure supplement 2C). Colocalization analysis indicates that treatment with phenformin does not change the amount of overlap between FARD-1::RFP and GFP::PTS1 relative to vehicle treated controls (Figure 2—figure supplement 2D). To confirm our earlier observation that suggests FARD-1 colocalization with lipid droplets, we used confocal imaging to assess the spatial distribution of an integrated FARD-1::RFP reporter *(fard-1 oe3)* in *C. elegans* fed C1-BODIPY-C12 to label lipid droplets (and treated with *glo-4* RNAi to remove BODIPY-positive lysosome-related organelles) (Hermann et al., 2005; Zhang et al., 2010a; Zhang et al., 2010b) We found that FARD-1::RFP fluorescence directly surrounds some, but not all, BODIPY-positive lipid droplets in the worm intestine (Figure 2—figure supplement 2E). However, as with peroxisomes, phenformin does not alter the number of lipid droplets that are surrounded by FARD-1 or its distribution around lipid droplets (data not shown.) Finally, FARD-1::RFP localizes into web-like structures in the *fard-1(oe3)* reporter that may represent endoplasmic reticulum versus another cellular tubular vesicular network (Figure 2—figure supplement 2F), and this localization is also not altered by biguanide treatment. Thus, the regulation of ether lipid biosynthesis does not appear to be via differential localization of FARD-1.

Expression of mRNAs encoding FARD-1, ACL-7, and ADS-1 are all decreased or unchanged in abundance upon treatment with biguanide via quantitative RT-PCR (Figure 2—figure supplement 2G—L), suggesting that ether lipids are not increased in phenformin treatment through a transcriptional mechanism. A parallel decrease in overall levels of FARD-1::RFP protein of *fard-1(oe1)* transgenics was seen with phenformin treatment (Figure 2—figure supplement 2M). These seemingly paradoxical data are likely consistent with post-translational negative feedback of ether lipids on the ether lipid biosynthetic pathway, as has been previously reported.

To affirm that the peroxisome is an essential site of ether lipid production in biguanide action, we disrupted peroxisomal protein targeting and examined phenformin-stimulated lifespan extension. Indeed, either *prx-5* or *prx-19* RNAi impair lifespan extension prompted by phenformin fully or partially, respectively (Figure 3A—B). PRX-5 is involved in protein import into the peroxisomal matrix and PRX-19 is involved in proper sorting of proteins for peroxisomal biogenesis. Thus, either disruption of ether lipid biosynthetic machinery or of a principal site of ether lipid biosynthesis impairs phenformin’s prolongevity benefit.

**Figure 3.**
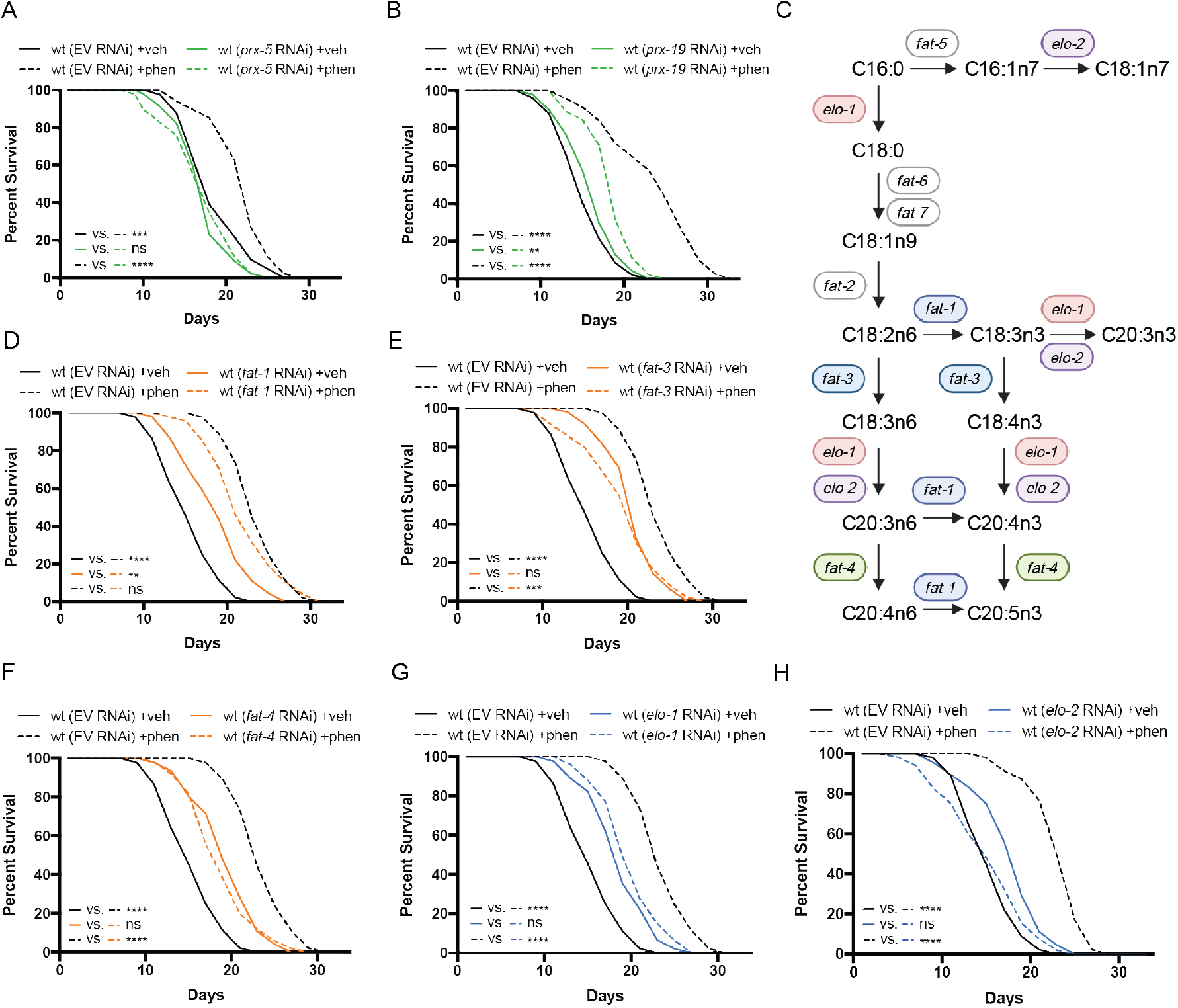
Peroxisomal protein import, fatty acid elongases, and fatty acid desaturases are required for the prolongevity effects of biguanides. (A-B) Knockdown of *prx-5* (A) and *prx-19* (B) by RNAi eliminates or significantly suppresses phenformin-mediated lifespan extension. (C) Schematic representation of the mono-(MUFA) and poly-unsaturated fatty acid (PUFA) synthesis pathway in *C. elegans* (adapted from (Watts, 2016)). (D-H) RNAi of three fatty acid desaturases (D-F) and two fatty acid elongases (G and H) involved in the synthesis of 18- and 20-carbon polyunsaturated fatty acids blunt phenformin-mediated lifespan extension in wild type worms. Colored symbols for *elo* and *fat* genes (vs. those in black and white) in Figure 3C indicates those that inhibit phenformin lifespan extension when knocked down by RNAi. For A, B and D-H, results are representative of 2-3 biological replicates. **, *P* < 0.01; ***, *P* < 0.001; ****, *P* < 0.0001 by log-rank analysis; see also Supplementary file 1 for tabular survival data and biological replicates.

### Fatty acid elongases and desaturases are positive effectors of biguanide-mediated lifespan extension

Most mature ether lipid species contain a fatty acid in the *sn-2* position linked by an ester bond (Dean and Lodhi, 2018). The majority of fatty acids conjugated in ether lipids are largely synthesized endogenously in *C. elegans* by fatty acid desaturases and fatty acid elongases (Perez and Van Gilst, 2008; Perez and Watts, 2021) (Figure 3C). Thus, we hypothesized that some of these desaturases and elongases may also contribute mechanistically to biguanide-mediated lifespan extension. Indeed, RNAi knockdown of three fatty acid desaturases and two fatty acid elongases in phenformin-treated *C. elegans* blunted phenformin-stimulated lifespan extension relative to empty vector controls (Figure 3D—H). Notably, these five genes all contribute to the production of fatty acids 18-20 carbons in length with three or more double bonds. Although knockdown of fatty acid desaturases and elongases in *C. elegans* results in inherent lifespan extension on vehicle relative to wild type controls on empty vector RNAi as has been previously reported (Horikawa et al., 2008; Shmookler Reis et al., 2011), these RNAi knockdown either largely (as in the case of *fat-1*, Figure 3D) or completely mitigate phenformin-driven lifespan extension (as in the case of *fat-3, fat-4, elo-1* and *elo-2*, Figure 3E—H). These results suggest the tantalizing possibility that fatty acid desaturases and elongases promote biguanide-mediated lifespan extension through contribution of long and polyunsaturated fatty acids to the synthesis of ether lipids, though a mechanistically distinct role is also possible.

### Genes involved in ether lipid biosynthesis are required in multiple longevity paradigms

Given the critical role of ether lipids in the response to biguanides, we hypothesized that these molecules may also play a broader role in diverse longevity paradigms involving metabolic or nutrient-sensing pathways. *C. elegans* mutant strains that exhibit 1) reduced mitochondrial function (*isp-1*), 2) disrupted mTORC1 signaling (*raga-1*), 3) abnormal pharyngeal pumping (*eat-2*), or 4) inhibition of insulin/insulin-like growth factor-1 signaling (*daf-2*), all result in extension of lifespan (Apfeld et al., 2004; Curtis et al., 2006; Schreiber et al., 2010; Senchuk et al., 2018). To determine whether requirement for the ether lipid biosynthetic machinery in aging generalizes to these other lifespan extension paradigms, we knocked down all three ether lipid biosynthetic enzymes by RNAi in wild type *C. elegans* and four long-lived genetic mutants: *raga-1, isp-1, eat-2*, and *daf-2*. Knockdown of *fard-1, acl-7*, and *ads-1* by RNAi results in suppression of lifespan extension in *isp-1, raga-1*, and *eat-2* mutants (Figure 4A—C). However, knockdown of ether lipid synthesis genes by RNAi did not impact lifespan extension in *daf-2* mutants (Figure 4—figure supplement 1A). Thus, the ether lipid biosynthetic machinery plays a broad role in lifespan extension, and, importantly, does not non-selectively shorten lifespan by making animals generally unfit.

**Figure 4.**
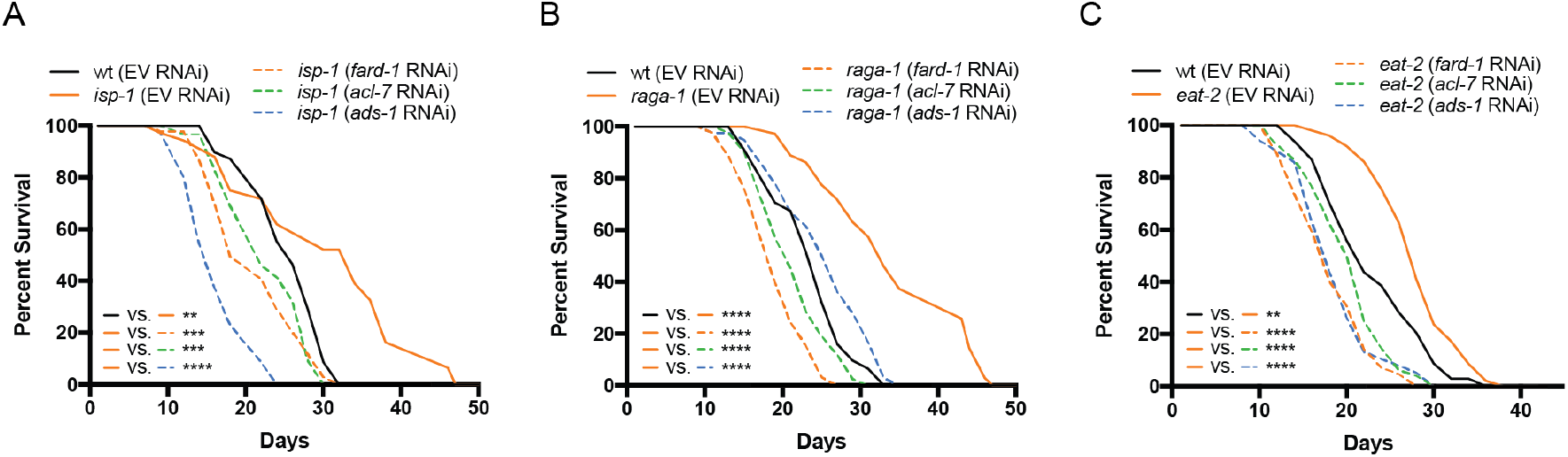
Genes involved in ether lipid biosynthesis are required for lifespan extension in multiple longevity paradigms. (A-C) *isp-1, raga-1*, and *eat-2* mutants display extended lifespan relative to wild type animals that is suppressed by RNAi knockdown of any of the three members of the ether lipid biosynthetic pathway. Results are representative of 3 biological replicates. **, *P* < 0.01; ***, *P* < 0.001; ****, *P* < 0.0001 by log-rank analysis. See also Figure 4—figure supplement 1 and Supplementary file 1 for tabular survival data and biological replicates.

### Overexpression of *fard-1* is sufficient to promote lifespan extension

To determine whether stimulation of ether lipid biosynthesis is sufficient to prompt lifespan extension, we tested the effect of overexpression (*oe*) of the sole *C. elegans* fatty acid reductase that synthesizes fatty alcohols for ether lipid biogenesis, *fard-1*, on lifespan. Strikingly, *fard-1(oe1)* alone significantly extends lifespan (Figure 5A). This result is similar in a second, independent *fard-1(oe2)* transgenic line (Figure 5B). To confirm that *fard-1(oe)* lifespan extension is dependent upon ether lipid biosynthesis, we knocked down *fard-1, acl-7*, and *ads-1* by RNAi in the *fard-1(oe1)* transgenic. As predicted, knockdown of three ether lipid biosynthetic enzymes leads to significant suppression of *fard-1(oe1)* lifespan extension (Figure 5C and (Figure 4—figure supplement 1B—C).

**Figure 5.**
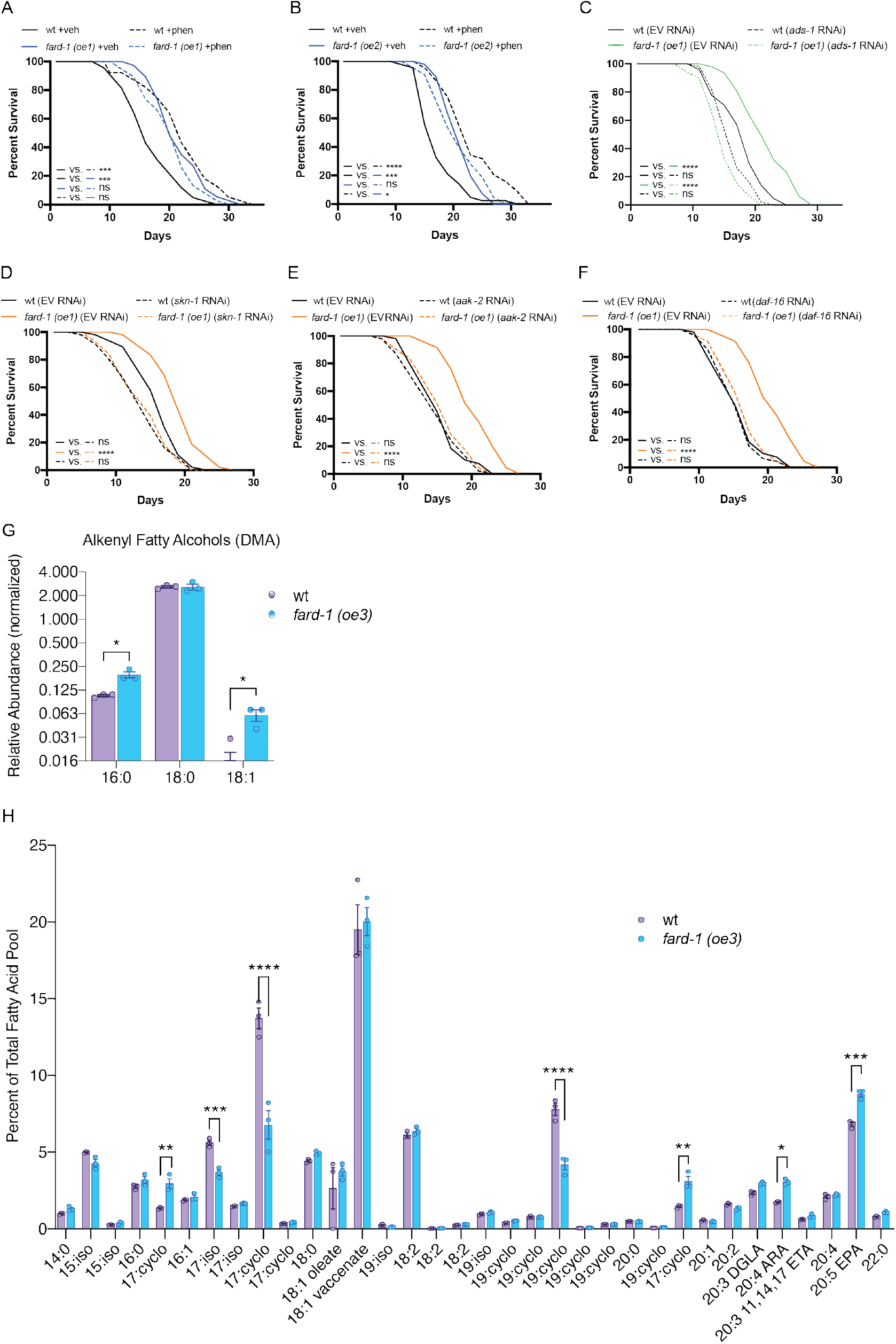
FARD-1 overexpression is sufficient to extend lifespan by modulating ether lipid synthesis. (A-B) Two, independently-generated *fard-1* overexpression *(fard-1 oe1 and fard-1 oe2)* transgenic strains exhibit lifespan extension that is not further extended by concomitant phenformin treatment. (C) RNAi knockdown of *ads-1* fully suppresses *fard-1(oe1)* lifespan extension, indicating that the *fard-1(oe)*-mediated lifespan extension is dependent upon ether lipid synthesis. (D-F) RNAi of *skn-1* (D), *aak-2* (E), and *daf-16* (F) suppress *fard-1(oe1)-*mediated lifespan extension. For *A*-*F*, results are representative of 2-3 biological replicates. *, *P* < 0.05; ***, *P* < 0.001; ****, *P* < 0.0001 by log-rank analysis. See also Figure 4—figure supplement 1 and Supplementary file 1 for tabular survival data and biological replicates. (G) Worms overexpressing integrated FARD-1 *(fard-1 oe3)* display a significant increase in 16:0 and 18:1 but not 18:0 alkenyl ether lipids by GC/MS. (H) Comparison of the total fatty acid pool indicates that the polyunsaturated fatty acids 20:4 arachidonic acid (ARA) and 20:5 eicosapentaenoic acid (EPA) are significantly increased in *fard-1* overexpressing *(fard-1 oe3)* worms vs. wild type animal, while several isomethyl (iso) and cyclopropyl (cyclo) fatty acids change in opposing directions. For *G-H*, n = 3 biological replicates. *, *P* < 0.05; **, *P* < 0.01; ***, *P* < 0.001; ****, *P* < 0.0001 by multiple t-tests (with multiple hypothesis correction by two-stage step-up method of Benjamini, Krieger, and Yekutieli).

To determine whether lifespan extension attributable to *fard-1(oe)* shares genetic dependencies with biguanide-mediated longevity, we independently knocked down *skn-1*/*Nrf2, aak-2/*AMPK and *daf-16*/*FoxO* by RNAi in a *fard-1(oe)* background. While *skn-1*/*Nrf2* and *aak-2/*AMPK have previously been demonstrated to be necessary for metformin-stimulated lifespan extension, *daf-16/FoxO* has not (Kenyon et al., 1993; Onken and Driscoll, 2010). Lifespan extension attributable to *fard-1(oe1)* is suppressed by these three gene knockdowns (Figure 5D—F), indicating that it is mechanistically similar, but not identical, to biguanide-mediated lifespan extension (Cabreiro et al., 2013; Onken and Driscoll, 2010). In aggregate, these results support the notion that ether lipids are an important requirement in multiple, diverse longevity paradigms, and further that *fard-1(oe)* promotes mechanistically-distinct lifespan extension in *C. elegans*.

In order to characterize shifts in ether lipids related to prolongevity effects, we performed comparative GC-MS-based fatty acid profiling of our integrated *fard-1(oe)* animals. Levels of 16:0 and 18:1 alkenyl ether lipids (indicated by DMAs on GC-MS analysis), are significantly increased in *fard-1(oe3)* transgenic animals versus wild type worms (Figure 5G). By comparison, 18:0 ether lipids were not increased, indicating that the ether lipid pool has both similarities and differences between *fard-1* overexpression and phenformin treatment. Echoing the analysis seen with phenformin treatment, few differences were found in a comparison of the relative abundance of fatty acids within the total lipid pool for *fard-1(oe3)* and wild type worms (Figure 5H). Those exhibiting increases in *fard-1(oe)* include the polyunsaturated fatty acids (PUFA) 20:4 arachidonate and 20:5 eicosapentaenoate (EPA). This suggests either that PUFAs play a mechanistic role in lifespan extension in *fard-1(oe)* or that they are increased as a consequence of longevity-promoting activity of ether-lipids.

### Ether lipids do not promote lifespan extension by modulating ferroptosis

Ether lipids have been reported to be protective against ferroptosis, an iron-dependent form of programmed cell death characterized by the accumulation of lipid peroxides (Perez et al., 2020; Zou et al., 2020). In order to determine whether ether lipids promote longevity downstream of biguanide action by modulating ferroptosis, we knocked down members of the glutathione peroxidase (GPX) family in animals overexpressing integrated *fard-1 (fard-1 oe3* and *fard-1 oe4)*, as has been previously reported to genetically facilitate lipid peroxidation and ferroptosis (Perez et al., 2020; Sakamoto et al., 2014) (Figure 5—figure supplement 1A—C). This analysis indicates that *gpx-1* (ortholog of human GPX4) RNAi leads to variable lifespan extension relative to wild type controls and exhibits non-additive lifespan extension with *fard-1(oe)* (Figure 5—figure supplement 1A). Neither *gpx-6* nor *gpx-7* knockdown impacts lifespan extension in *fard-1(oe)* animals (Figure 5—figure supplement 1B—C). Further, GPX family RNAi do not negatively impact lifespan extension reproducibly downstream of phenformin (Figure 5—figure supplement 1D—F). We conclude that genetic triggers that induce ferroptosis do not impact phenformin-prompted or *fard-1(oe)* lifespan extension, and thus it is unlikely that either extend lifespan by suppressing ferroptosis.

### The ether lipid biosynthetic machinery operates upstream of the stress responsive factor, *skn-1/Nrf2*, to enable lifespan extension in response to biguanides

We noted when analyzing FARD-1 protein localization that somatic lipid droplets are generally less numerous in BODIPY-stained, phenformin-treated animals vs. vehicle. Indeed, quantitative analysis indicates that intestinal lipid droplets are significantly less numerous following phenformin treatment (in *glo-4* RNAi-treated FARD-1::RFP transgenics *(fard-1 oe3)* fed C1-BODIPY-C12 to label lipid droplets, Figure 6A). We previously reported that gain-of-function mutations in the nutrient- and stress-responsive transcription factor *skn-1/Nrf2* prompt age-dependent, somatic depletion of fat (Asdf) (Lynn et al., 2015; Nhan et al., 2019). This, together with early adult decreases in lipid droplet numbers, suggested to us that phenformin may prompt longevity by activating metabolic stress defenses in a *skn-1-*dependent manner. Strikingly, we found that phenformin treatment produces Asdf at day three of adulthood, quantitatively analogous to and non-additive with *skn-1* gain-of-function mutants (Figure 6B—C). Compellingly, loss of function mutations in any of the three ether lipid biosynthetic genes completely prevent the phenformin-mediated Asdf phenotype (Figure 6B—C). Together with the observation that promotion of lifespan extension by both phenformin and *fard-1(oe)* require *skn-1*, these data suggest that biguanides activate an ether lipid-*skn-1* signaling relay to drive longevity-associated metabolic shifts.

**Figure 6.**
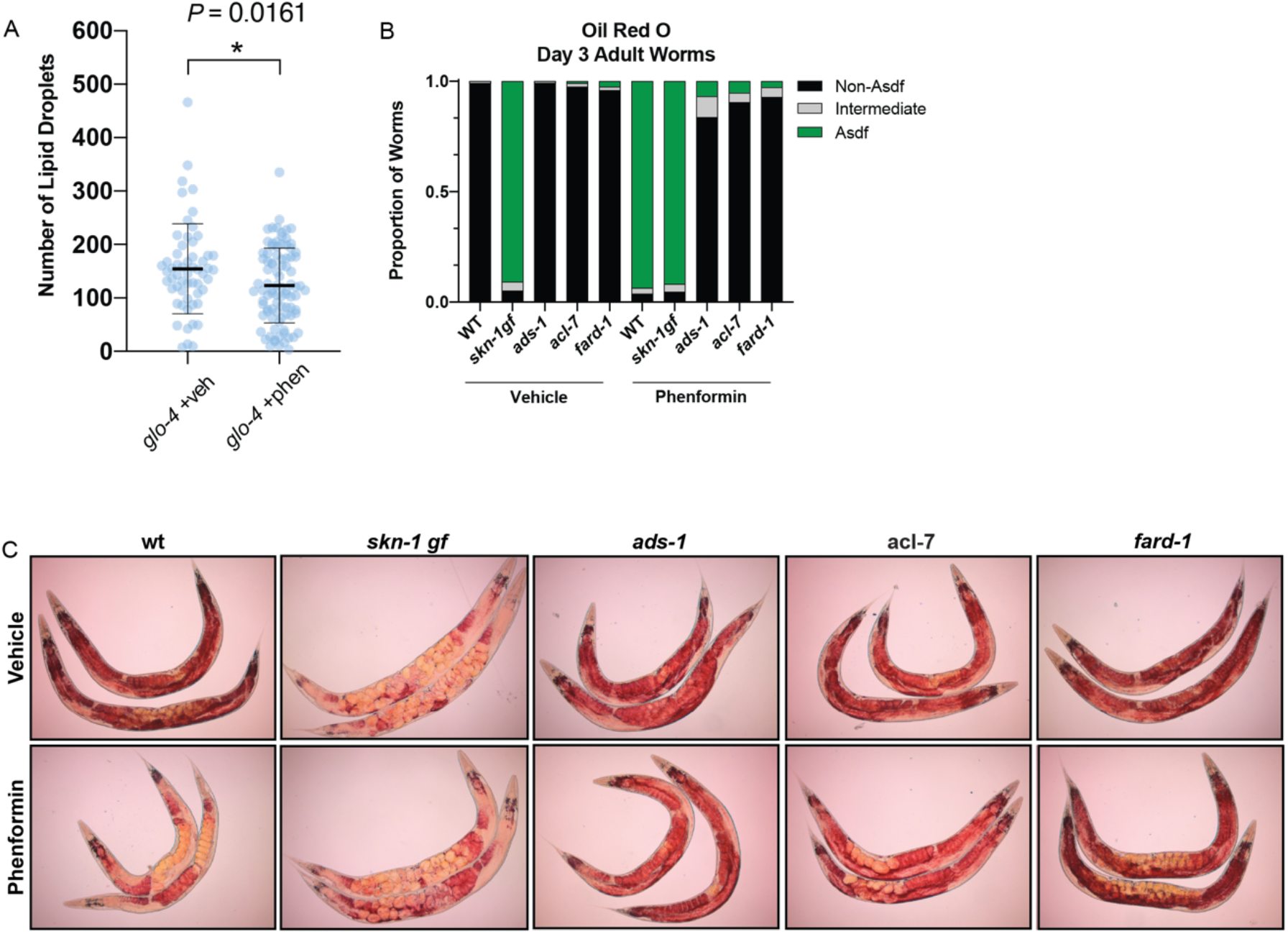
Phenformin modulates systemic lipid metabolism through an ether lipid-*skn-1* signaling relay. (A) The number of intestinal, C1-BODIPY-C12 labeled lipid droplets are significantly lower in phenformin treated animals versus vehicle (FARD-1::RFP reporter transgenic *(fard-1 oe3)* worms are also treated with *glo-4* RNAi to remove BODIPY-positive lysosome-related organelles). n=2 biological replicates. *, *P* < 0.05 by unpaired t-test. (B-C) Oil-red-O staining of day 3 adult phenformin treated wild type animals indicates that drug treatment leads to age-dependent somatic depletion of fat (Asdf), as previously reported for *skn-1* gain of function mutants (*skn-1 gf)*, suggesting that phenformin activates Asdf downstream of *skn-1*. Quantification (B) indicates that the proportion of Asdf animals is non additively increased by phenformin treatment in a *skn-1gf* mutant, and that phenformin is no longer able to activate Asdf in 3, independent ether lipid deficient mutants in *ads-1, acl-7*, and *fard-1*. For *B-C*, n=3 biological replicates.

## Discussion

In an unbiased RNAi screen of ∼1000 metabolic genes, we identified ether lipid biosynthesis as critical to the longevity-promoting effects of metformin in *C. elegans*. Our results show that the biguanides metformin and phenformin promote lifespan extension by stimulating biogenesis of ether lipids, prompting longevity-promoting, metabolic stress defenses mediated by *skn-1*. The broad importance of ether lipids is demonstrated by their requirement in multiple, diverse paradigms of lifespan extension. Our findings also indicate that ether lipid modulation through overexpression of *fard-1* is also sufficient to promote longevity. Thus, ether lipids form a heretofore unappreciated lynchpin of lifespan modulation, and are sufficient to support healthy aging through multiple, central longevity effectors, including *skn-1*.

Differences in ether lipid abundance and composition are correlated with diseases of aging. The uniform lethality associated with human genetic ether lipid deficiency, as in the case of patients diagnosed with RCDP and Zellweger syndrome, has made it difficult to study the role of ether lipids in aging and aging-associated diseases (Braverman et al., 1997; Itzkovitz et al., 2012; Motley et al., 1997; Purdue et al., 1997). Nonetheless, observational studies demonstrate decreases in certain plasmalogen species in Alzheimer’s Disease, suggesting a probable link between ether lipids and aging-related pathologies (Goodenowe et al., 2007; Grimm et al., 2011; Han et al., 2001). Ether lipids have conflicting roles in cancer; while loss of the ether lipid biosynthetic machinery profits cancer cell survival by enhancing resistance to ferroptosis (Zou et al., 2020), in other contexts, ether lipid deficiency results in impaired pathogenicity in various human cancer cells (Benjamin et al., 2013; Perez et al., 2020). Cancer cells generally have higher levels of ether lipids compared to normal cells, leading others to suggest that ether lipids confer pro-survival benefit (Albert and Anderson, 1977; Benjamin et al., 2013; Snyder and Wood, 1969). However, certain ether lipid species have also been reported to have anti-tumor properties (Arthur and Bittman, 2014; Jaffrès et al., 2016). Thus, in line with the results we present here, it is critical to understand ether lipids in context. Future work will need to focus on the impact of specific ether lipid species rather than the whole class *en masse* to understand which may play a beneficial versus detrimental role in health.

Studies in long-lived animal models suggest there is an association between ether lipid content and animal longevity, such as in the naked mole-rat (*Heterocephalus glaber*) (Mitchell et al., 2007) and the mud clam *Arctica islandica* (Munro and Blier, 2012). Higher plasmalogen levels in naked mole-rat tissues versus mice are speculated to contribute to protection of cellular membranes via a reduction of oxidative stress (Mitchell et al., 2007). Similarly, exceptionally long-lived humans harbor higher levels of phosphatidylcholine-derived, short chained alkyl ether lipids and a lower levels of phosphatidylethanolamine-derived longer chained plasmalogens (Pradas et al., 2019), but these associations are of unclear functional significance. Although it is clear that ether lipid deficiency in *C. elegans* prevents longevity downstream of mitochondrial electron transport chain dysfunction, mTOR deficiency, caloric restriction, and biguanides alike, the precise lipid(s) conferring this activity remains unknown. Each of these longevity paradigms have features of nutrient deficiency, energy stress, or nutrient sensing, so it is possible that ether lipids are at least part of the common effector arm conferring benefit in aging to various forms of metabolic stress.

Our results suggest that unsaturated fatty acids and phosphatidylethanolamine ether lipids are essential to the health promoting effects of biguanides. Although we see major shifts in abundance of alkenyl ether lipids, genetic evidence of necessity of ether lipids, and requirement for the synthesis of mono- and poly-unsaturated fatty acids in biguanide-induced longevity, determination of the specific lipids necessary for promoting healthy aging awaits the ability to modulate the level of specific ether lipids. Additionally, disruption of ether lipid biosynthesis has been shown to increase the proportion of stearate (18:0) and other saturated fatty acids (Shi et al., 2016). Thus, at this time, we cannot rule out the possibility that biguanide-stimulated alterations in ether lipid biosynthesis serves to divert accumulation of lipid species that are detrimental to lifespan, for instance, saturated fatty acids. Nonetheless, in light of our finding that ether lipids prompt metabolic stress defenses, this alternative mechanism is less likely. Definitive proof will require a deeper understanding of the regulation of specific steps dictating the synthesis and modification of ether lipids of different fatty alcohol and fatty acid composition.

Based upon our findings, ether lipid synthesis is likely to be regulated post-translationally by biguanide treatment. The demonstrated increase in plasmalogens and specific ether lipids are both consistent with increases in activity of the ether lipid biosynthetic machinery. While we do not understand the mechanism for the increased activity of ether lipid synthesizing enzymes, the decreases in mRNAs for *acl-7, ads-1*, and *fard-1* and protein for FARD-1 invokes negative feedback consistent with previous work showing that higher levels of ether lipids promotes proteasomal degradation of peroxisomal Far1 protein (Honsho et al., 2010). Co-localization of the fatty alcohol reductase, FARD-1, with both peroxisomes and lipid droplets is similarly not impacted by biguanides. Further investigation into the precise molecular interactions between FARD-1 protein and other organelles will be required to further understand how FARD-1 and the other ether lipid biosynthetic enzymes are regulated by biguanides and in aging.

Strikingly, our data demonstrate for the first time that ether lipids are required for phenformin to activate metabolic defenses downstream of the stress- and metabolism-responsive transcription factor *skn-1*/NRF2. Phenformin drives age-dependent somatic depletion of fat (Asdf), a phenotype we previously reported upon genetic activation of *skn-1* (Lynn et al., 2015; Nhan et al., 2019). Based upon the work of others and our own unpublished work, biguanides do not potentially stimulate canonical *skn-1* antioxidant defenses such as *gst-4* expression (Cabreiro et al., 2013; Onken and Driscoll, 2010). We suggest that *skn-1* is uniquely required for metabolic stress defenses downstream of metformin such as Asdf rather than canonical oxidative or proteostatic defenses. The requirement for ether lipids in Asdf activation by phenformin confirm that this class of lipids plays a heretofore unappreciated role in a distinct form of *skn-1* activation mimicked by genetic forms of *skn-1* activation we have previously reported (Lynn et al., 2015; Nhan et al., 2019).

In aggregate, data presented here indicate that ether lipid biosynthesis plays a broader role in aging than previously described. The necessity of the ether lipid machinery in metformin- and phenformin-stimulated lifespan extension and in multiple longevity paradigms indicates that ether lipids serve as a lynchpin through which lifespan is modulated (Figure 7A—B). Our demonstration that overexpression of FARD-1 alone results in lifespan extension provides an exciting opportunity to identify ether lipids that promote health and the effector mechanisms through which they act. Finally, these results support the exciting possibility that modulation of ether lipids pharmacologically or even dietarily may provide a new potential therapeutic target in aging and aging-related diseases.

**Figure 7.**
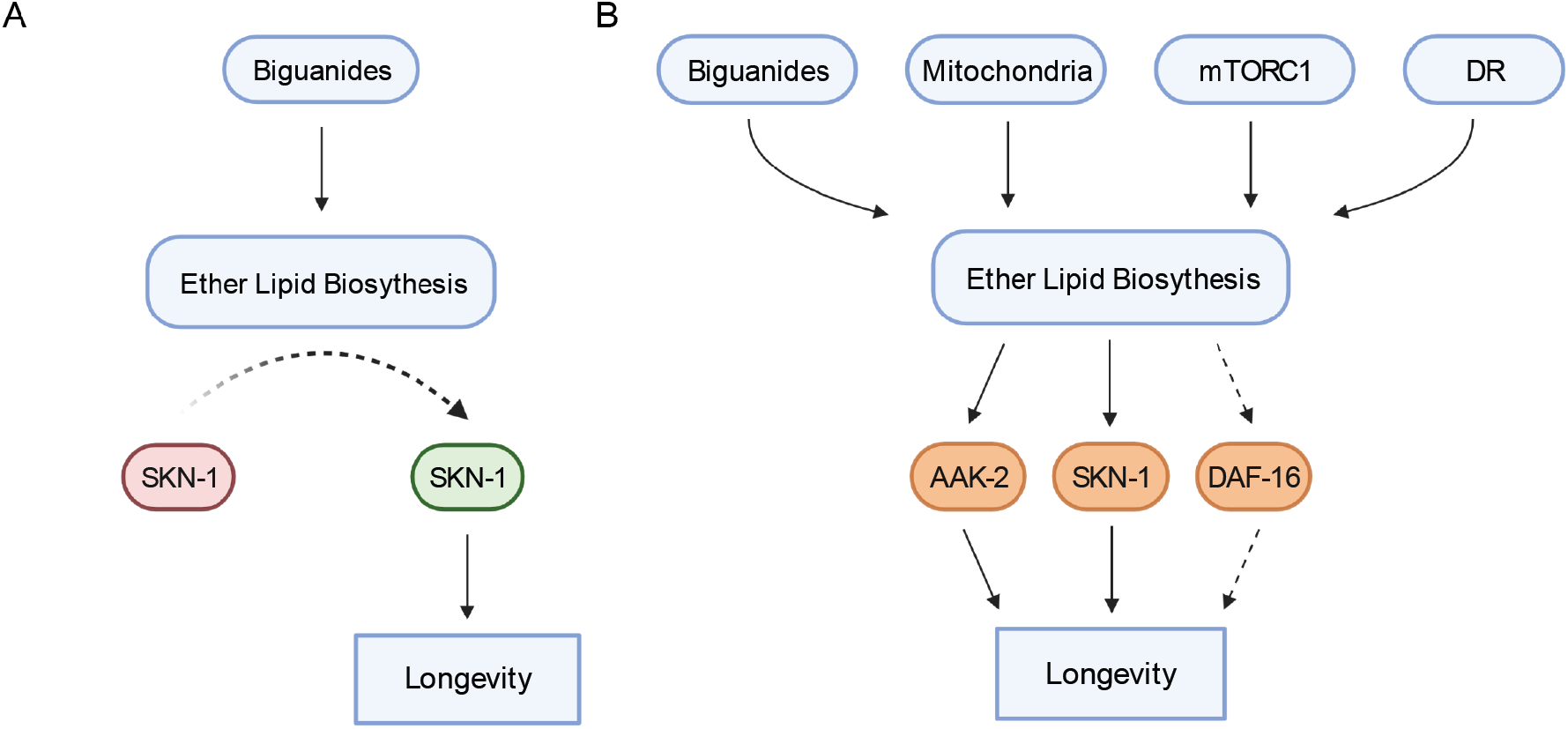
Schematic representation for the role of the ether lipid biosynthetic machinery in multiple prolongevity paradigms. (A) Model of ether lipid action in biguanide-prompted lifespan extension. Activation of ether lipid biosynthesis leads to longevity-promoting activity of metabolic stress defenses downstream of the transcription factor *skn-1*. (B) Model portraying a broader than previously appreciated role of ether lipids in longevity downstream of biguanides, mitochondrial electron transport inhibition, mTORC1 inhibition, and dietary restriction (DR).

## Materials and Methods

### *C. elegans* genetics

Strains were maintained at 20°C grown on E. coli OP50-1 (RRID: WB-STRAIN:WBStrain00041971) for all experiments unless otherwise indicated. The following strains were used in this study: N2 (wild type strain, RRID: WB-STRAIN:WBStrain00000001), BX275 *fard-1(wa28)* (RRID: WB-STRAIN:WBStrain00004025), BX259 *acl-7(wa20*) (RRID: WS-STRAIN: WBStrain00004024), BX10 *ads-1(wa3*) (RRID: WB-STRAIN:WBStrain00004007), CB1370 *daf-2(e1370*) (RRID: WB-STRAIN: WBStrain00004309), MQ989 *isp-1(qm150*) (RRID: WB-STRAIN: WBStrain00026672), VC533 *raga-1(ok701*) (RRID: WB-STRAIN: WBStrain00035849), DA465 *eat-2(da465*) (RRID:WB-STRAIN: WBStrain00005463), MGH48 *mgIs43*[*ges-1p*::GFP::PTS1], SPC168 *skn-1*(*lax188*) (*skn-1gf*, RRID: WB-STRAIN: WBStrain00034420). For *fard-1* overexpression, the following strains were generated: MGH471 *alxEx122*[*fard-1p*::FARD-1::mRFP::HA unc-54 3’UTR *myo-2p*::GFP] *(fard-1 oe1)*, MGH472 *alxEx135*[*fard-1p*::FARD-1::mRFP::HA unc-54 3’UTR *myo-2p*::GFP] *(fard-1 oe2)*, MGH605 *alxIs45*[*fard-1p*::FARD-1::mRFP::HA::unc-54 3’UTR *myo-2p*::GFP] *(fard-1 oe3)*, and MGH606 *alxIs46*[*fard-1p*::FARD-1::mRFP::HA::unc-54 3’UTR *myo-2p*::GFP] *(fard-1 oe4)*. All strains for *fard-1* overexpression were backcrossed 8x to N2 Bristol. For colocalization analysis with peroxisomally targeted GFP, we crossed MGH48 and MGH471 to generate the strain: MGH607 *mgIs43*[*ges-1p*::GFP::PTS1]; *alxEx122*[*fard-1p*::FARD-1::mRFP::HA::unc-54 3’UTR *myo-2p*::GFP] (noted in text as GFP::PTS1; FARD-1::RFP).

### Generation of *fard-1 C. elegans* transgenic lines

For FARD-1 expression, the entire genomic sequence of the *fard-1* locus (3659 bp), including introns and exons, plus 4910 bp of promoter were amplified and cloned into a modified Fire vector driving *fard-1* fused to mRFP and a HA epitope tag at the C-terminus. The following cloning primers were used:

F: 5**’**-TGCATGCCTGCAGGTCGACTTTGACAAAAGTTCTGTTGCCG-3**’** and R:

5’-TTTGGGTCCTTTGGCCAATCGCTTTTTTGAAGATACCGAGAATAATCC-3’.

The FARD-1 overexpression construct was injected at 10 ng/μL (*alxEx122*) and 18 ng/μL (*alxEx135*) into the gonad of wild type adult animals with salmon sperm DNA as a carrier and 1.5 ng/μL *myo-2p*::GFP as a co-injection marker. *alxEx122* was subsequently integrated by UV irradiation and 8x backcrossed to N2 Bristol to obtain MGH605 and MGH606.

### RNA interference (RNAi) assays

RNAi clones were isolated from a genome-wide *E. coli* RNAi library (generated in strain HT115(DE3), RRID: WB-STRAIN:WBStrain00041079), sequence verified, and fed to animals as described (Kamath and Ahringer, 2003). RNAi feeding plates (6 cm) were prepared using a standard NGM recipe with 5 mM isopropyl-B–D-thiogalactopyranoside and 200 μg/ml carbenicillin. RNAi clones were grown for 15 hours in Luria Broth (LB) containing 100 μg/ml carbenicillin with shaking at 37°C. The stationary phase culture was then collected, concentrated through centrifugation, the supernatant was discarded, and the pellet was resuspended in LB to 20% of the original culture volume; 250 μl of each RNAi clone concentrate was added to RNAi plates and allowed to dry at least 24 hours prior to adding biguanide. Drug treatment was added to seeded RNAi plates and allowed to dry at least 3 hours before adding worms.

### Longevity assays

Lifespan analysis was conducted at 20°C, as previously described (Soukas et al., 2009). Briefly, synchronized L1 animals were seeded onto NGM (for mutant treatment) or RNAi plates (for RNAi) and allowed to grow until the L4 to YA transition stage. On day 0 of adulthood as indexed in the figure legend, ∼50-60 L4/YA worms per plate (unless otherwise noted) were transferred onto fresh NGM or RNAi plates. These NGM and RNAi plates were supplemented with 30 μM and 100 μM 5-fluorodeoxyuridine (FUdR) to suppress progeny production, respectively. For biguanide treatment, about ∼55-60 synchronized L1 animals (unless otherwise noted) were seeded onto plates containing 50 mM metformin or 4.5 mM phenformin. Based upon power calculation for log-rank analysis, minimum N of 50 (per group) was chosen to satisfy α = 0.05, β = 0.2, and effect size = 20% difference in lifespan (Petrascheck and Miller, 2017). At the L4/YA stage, these worms were transferred to plates containing biguanide treatment and FUdR for the remainder of their life. Dead worms were counted every other day, and scoring investigators were blinded as to the experimental group/treatment until the conclusion of each experiment. Statistical analysis was performed with online OASIS2 resources (Han et al., 2016).

### Body Size Determination of *C. elegans*

We measured worm body size in response to biguanide treatment by imaging as previously described (Wu et al., 2016). Egg prep synchronized wild type worms were treated with empty vector (L4440) or ether lipid biosynthesis machinery RNAi and treated with vehicle (ddH_2_O) or 160mM metformin. After ∼65-70 hours, worms were transferred into a 96 well plate, washed 3x with M9, and paralyzed in M9 buffer with 1 mg/ml levamisole (L9756-10G, Sigma-Aldrich). Once immobilized, brightfield imaging was performed at 5X magnification on a Leica DM6000 microscope within 5 minutes of transferring to a 96 well Teflon imaging slide. We determined the maximal, longitudinal cross-sectional area of the imaged worms by using MetaMorph software for a minimum of ∼80 animals per condition in each experiment. Results of a single experiment is shown. Each experiment was performed at least twice, and results were consistent between experiments.

### GC/MS lipidomics

Lipid extraction and GC/MS of extracted, acid-methanol-derivatized lipids was performed as described previously (Pino and Soukas, 2020; Pino et al., 2013). Briefly, 5000 synchronous mid-L4 animals were sonicated with a probe sonicator on high intensity in a microfuge tube in 100-250 microliters total volume. Following sonication, lipids were extracted in 3:1 methanol: methylene chloride following the addition of acetyl chloride in sealed borosilicate glass tubes, which were then incubated in a 75°C water bath for 1 hour. Derivatized fatty acids and fatty alcohols were neutralized with 7% potassium carbonate, extracted with hexane, and washed with acetonitrile prior to evaporation under nitrogen. Lipids were resuspended in 200 microliters of hexane and analyzed on an Agilent GC/MS equipped with a Supelcowax-10 column as previously described (Pino and Soukas, 2020). Fatty acids and alcohols are indicated as the normalized peak area of the total of derivatized fatty acids and alcohols detected in the sample. Based upon power calculation for pairwise comparison, a minimum N of 3 biological replicates (per group) was chosen to satisfy α = 0.05, β = 0.2, and effect size = 50% with σ = 20%. Analyses were blinded to the investigator conducting the experiment and mass spectrometry calculations until the conclusion of each experiment when aggregate statistics were computed.

### LC/MS-MS lipidomics

Wild type, *fard-1, acl-7*, and *ads-1* worm mutants were collected using conditions that enabled our reported longevity phenotypes. Briefly, collection for LC/MS-MS processing comprised of 3 replicates of these 4 strains that were independently treated with vehicle (ddH_2_O) and 4.5mM phenformin on 10cm NGM plates. Based upon power calculations, as for GC/MS, a minimum N of 3 biological replicates (per group) was chosen to satisfy α = 0.05, β = 0.2, and effect size = 50% with σ = 20%, though the power is only expected to hold for the first significant difference detected. Analyses were blinded to the investigator conducting the experiment and mass spectrometry calculations until the conclusion of each experiment when aggregate statistics were computed. A total of ∼6,000 animals (2 × 10cmM plates, 3,000 worms per plate) were utilized per sample. These worms were washed with M9 (4x), concentrated into 200 μL of M9, and then flash frozen with liquid nitrogen in 1.5mL Eppendorf microcentrifuge tubes. Worm pellets were transferred to 2 mL impact resistant homogenization tubes containing 300 mg of 1 mm zirconium beads and 1 mL of 90:10 ethanol:water. Using a Precellys 24 tissue homogenizer, samples were homogenized in three 10 second cycles at 6400 Hz followed by 2 minutes of sonication. Samples were then placed at -20 °C for one hour to facilitate protein precipitation. Samples were transferred to 1.5 mL microfuge tubes and centrifuged at 14,000 g for 10 minutes at 4 °C. After centrifugation, 120 µL of supernatant was dried *in vacuo* and resuspended in 120 µL of 80:20 methanol:water containing internal standards 1 ng/µL CUDA and 1 ng/µL MAPCHO-12-d38. Lipidomic data was acquired by injecting 20 µL of sample onto a Phenomenex Kinetex F5 2.6 µm (2.1 × 100 mm) column at 40 °C and flowing at 0.35 mL/min. Metabolites were eluted using (A) water containing 0.1% formic acid and (B) acetonitrile:isopropanol (50:50) containing 0.1% formic acid using the following gradient: 0% B from 0-1 min, 0-50% B from 1-6 mins, 50-100% B from 6 to 17 minutes and 100% B hold from 17-20 mins. Compounds were detected using a Thermo Scientific QExactive Orbitrap mass spectrometer equipped with a heated electrospray ionization (HESI) source operating in positive and negative ion mode with the following source parameters: sheath gas flow of 40 units, aux gas flow of 15 units, sweep gas flow of 2 units, spray voltage of +/-3.5 kV, capillary temperature of 265°C, aux gas temp of 350°C, S-lens RF at 45. Data was collected using an MS1 scan event followed by 4 DDA scan events using an isolation window of 1.0 m/z and a normalized collision energy of 30 arbitrary units. For MS1 scan events, scan range of m/z 100-1500, mass resolution of 17.5k, AGC of 1e^6^ and inject time of 50 ms was used. For tandem MS acquisition, mass resolution of 17.5 k, AGC 5e^5^ and inject time of 80 ms was used. Data was collected using Thermo Xcalibur software (version 4.1.31.9) and analyzed using Thermo QualBrowser (version 4.1.31.9) as well as MZmine 2.36.

### Statistical analysis of metabolomics data

All visualization and significance testing of metabolomics was conducted using the MetaboAnalyst 5.0 package (Pang et al., 2021). Mass integration values for 9,192 compounds were extracted from full-scan LC-MS/MS measurements of L4 to young adult (YA) transition wild type (N2 Bristol), *ads-1(wa3), acl-7(wa20)*, and *fard-1(wa28)* animals treated from L1 hatch with vehicle, 4.5 mM phenformin, or 50mM metformin. Missing and zero values in the data matrix were imputed via replacement with 1/5th of the minimum positive value for each variable. Abundance values were subsequently filtered based on interquartile range (reducing the compound list to the 2500 most variable compounds), and log_10_ transformed. Quantile normalization was then performed, followed with division by the standard deviation of each variable (auto-scaling). Normalized abundance values for were then extracted based upon MS/MS signatures for phosphatidylethanolamine ether lipids and assessed for statistical significance via one-way ANOVA followed by false discovery rate (FDR) control using the Benjamini-Hochberg (BH) method (Benjamini and Hochberg, 1995). Post-hoc testing was then performed using Fisher’s LSD to evaluate pairwise comparison significance. Metabolites were considered differentially abundant in any one condition with an FDR controlled P value < 0.05. The top 25 metabolites across treatment (ranked by ANOVA f statistic and FDR value) were visualized using a heatmap of Euclidean distance measurements, with Ward clustering of samples and normalized compound abundances included. All mass integration values for identified phosphatidylethanolamine containing ether lipids, normalized abundance values, and log-transformed, normalized abundance values, are included in this manuscript as Supplementary file 2.

### Quantitative RT-PCR

To assess changes in mRNA levels of *fard-1, acl-7*, and *ads-1* in response to biguanide treatment, we used quantitative RT-PCR as previously described (Wu et al., 2016). Briefly, synchronized wild type (N2) L1 animals were seeded onto OP50, NGM plates containing vehicle (ddH_2_O), 50 mM metformin, or 4.5 mM phenformin. ∼1600 worms were collected from 4 6cm plates per replicate, per condition (with no more than 400 worms seeded per plate to prevent overcrowding). n = 3 biological replicates. Worms were collected at the L4 to YA transition using M9 buffer and washed an additional 3X, allowing worms to settle by gravity between washes. Total RNA was extracted using TRIzol. Reverse transcription was performed with the Quantitect reverse transcription kit (Qiagen). qRT-PCR was conducted in triplicate using Quantitect SYBR Green PCR reagent (Qiagen) following manufacturer instructions on a Bio-Rad CFX96 Real-Time PCR system (Bio-Rad). If not processed immediately, worms were flash frozen in liquid nitrogen and kept in −80 °C until RNA preparation. The sequences for primer sets used in *C. elegans* are:

*act-1*:

F: 5’-TGCTGATCGTATGCAGAAGG-3’ and

R: 5’-TAGATCCTCCGATCCAGACG-3’

*fard-1*:

F: 5’-ACAAGTCACCAATGGCTCCAC-3’ and

R: 5’-GCTTTGGTCAGAGTGTAGGTG-3’

*acl-7*:

F: 5’-GTTTATGGCTGGCGTGTTG-3’ and

R: 5’-CGGAGAAGACAGCCCAGTAG-3’

*ads-1*:

F: 5’-GCGATTAACAAGGACGGACA-3’ and

R: 5’-CGATGCCCAAGTAGTTCTCG-3’.

Expression levels of tested genes were presented as normalized fold changes to the mRNA abundance of *act-1* for *C. elegans* by the ΔΔCt method.

### FARD-1 overexpression reporter fluorescence intensity analysis

To assess changes in levels of fluorescent FARD-1 protein in response to biguanide treatment, we used the strain MGH471 *alxEx122*[*fard-1p*::FARD-1::mRFP::HA unc-54 3’UTR *myo-2p*::GFP] *(fard-1 oe1)*. In brief, egg prep synchronized L1 FARD-1::RFP transgenic worms were treated with vehicle (ddH_2_O) or 4.5 mM phenformin, paralyzed with 1 mg/ml of levamisole, and then imaged in 96-well format with a Leica DM6000 microscope outfitted with a mCherry filter set and MMAF software. These imaging experiments were carried out in biological triplicate with ∼10 animals imaged per replicate. Images were qualitatively assessed to obtain conclusions and results were consistent between independent replicates.

### Colocalization analysis of FARD-1::RFP and peroxisomally targeted GFP

Colocalization of GFP and RFP expression in vehicle or phenformin treated MGH607 was performed by Coloc2 (Fiji) on images taken on a Leica Thunder microscopy system. Since FARD-1::RFP in MGH607 is exogenously expressed, we performed 3 hour egg lays with ∼30 gravid hermaphrodites expressing both GFP::PTS1 and FARD-1::RFP to synchronize L1s. The eggs were treated with vehicle (ddH_2_O) or 4.5mM phenformin immediately after gravid hermaphrodites were removed, dried in a laminar flow hood, and allowed to incubate at 20°C until the worms were young adult/early day 1 adults. To prepare for imaging, only worms expressing both GFP::PTS1 and FARD-1::RFP were picked onto slides containing dried 2% agar pads, immobilized in ∼5 μL of 2.5mM levamisole solution and covered with a cover slip. Images of the upper, mid, and lower intestine were taken for 30 individual worms per condition (15 worms per replicate for 2 biological replicates). We generated Pearson’s r values to assess the extent to which intestinal RFP and GFP overlap in each region of all samples. All Pearson’s r values were combined to generate 4 individual averages (1 per condition) to perform an unpaired t-test.

### Lipid droplet analysis

The strain MGH605 *alxIs45*[*fard-1p*::FARD-1::mRFP ::HA::unc-54 3’UTR *myo-2p*::GFP] *(fard-1 oe3)* was used for this analysis. Preparation of worms for imaging was similar to our longevity assays but modified to incorporate staining of lipid droplets. Briefly, 6cm RNAi plates were seeded with 250μL bacteria expressing *glo-4* RNAi [5X] and allowed to incubate for 24 hours at 20°C. 1μM of green C1-BODIPY-C12 (D-3823, Invitrogen) diluted in 100μL 1X phosphate buffer saline (PBS, pH 7.2) was then added to the RNAi bacteria lawn as in (Soukas et al., 2009).The plates were immediately dried in a dark laminar flow hood, wrapped in aluminum foil to prevent photobleaching, and allowed to incubate at 20°C for 24-48 hours. These plates were treated with vehicle (ddH_2_O) or 4.5mM phenformin as mentioned previously (while kept away from light.) Egg prep synchronized worms were dropped onto plates and grown to day 1 adult stage. To prepare for confocal imaging, animals were rapidly picked onto slides containing dried 2% agar pads, immobilized in ∼5 μL of 2.5mM levamisole solution and covered with a cover slip. Lipid droplets were imaged by Zeiss LSM 800 Airyscan within 5 minutes of slide placement. Z-stacked images were obtained for the intestine near the tail end of 14 *glo-4*, vehicle treated and 19 *glo-4*, phenformin treated worms (2 biological replicates per condition). 5 planes were extracted (planes 1,2,4,5, and 9) using ImageJ for all samples. For lipid droplet counting, quantification was performed using CellProfiler 4.2.1 (Stirling et al., 2021) where lipid droplets were identified as primary objects. The min/max range for typical object diameters was 3-67 pixels, and those objects outside of the diameter range were discarded. Planes were excluded entirely if the pipeline did not accurately capture individual lipid droplets for the vast majority of objects.

### Oil Red O Staining

Oil red O (ORO) fat staining was conducted as outlined in (Stuhr et al., 2022), In brief, worms were egg prepped and allowed to hatch overnight for a synchronous L1 population. The next day, worms were dropped onto plates seeded with bacteria with or without phenformin and raised to 120 hours (day 3 adult stage). Worms were washed off plates with PBST, then rocked for 3 min in 40% isopropyl alcohol before being pelleted and treated with ORO in diH2O for 2 hours. Worms were pelleted after 2 hours and washed in PBST for 30 min before being imaged at 5x magnification with the DIC filter on the Zeiss Axio Imager Erc color camera.

### Asdf Quantification

ORO-stained worms were placed on glass slides and a coverslip was placed over the sample. Worms were scored, as previously described (Stuhr et al., 2022). Worms were scored and images were taken with the Zeiss Axio Imager Erc color camera at 5x magnification. Fat levels of worms were placed into three categories: non-Asdf, intermediate, and Asdf. Non-Asdf worms display no loss of fat and are stained dark red throughout most of the body (somatic and germ cells). Intermediate worms display significant fat loss from the somatic tissues, with portions of the intestine being clear, but ORO-stained fat deposits are still visible (somatic < germ cells). Asdf worms had most, if not all, observable somatic fat deposits depleted (germ cells only).

### Quantification and statistical analysis

Unless otherwise indicated, the statistical differences between control and experimental groups were determined by two-tailed students *t*-test (two groups), one-way ANOVA (more than two groups), or two-way ANOVA (two independent experimental variables), with corrected *P* values < 0.05 considered significant. Analyses conducting more than two comparisons were always corrected for multiple hypothesis testing. The log rank test was used to determine significance in lifespan analyses using online OASIS2.

## Acknowledgments

We thank Talia Hart, Dr. Gary Ruvkun, Dr. Eric Greer, and Dr. Keith Blackwell for discussions and constructive criticisms. This work was funded by NIH/NIA Grants R01AG058259 and R01AG69677 (to A.A.S.) and R01AG058610 (to S.P.C.), by the Weissman Family MGH Research Scholar Award (to A.A.S.), by a NSF GRFP Award 1000253984 (to L.C.), and by NIH/NIAID R01AI130289 (to R.P.W.), and by IRACDA NIH Grant K12GM106996 (to L.C.). Thanks to the University of Southern California and Buck Institute Nathan Shock Center (P30AG068345) for providing core services and support. Thanks to the NIH/NIDDK-funded NORC of Harvard (P30DK040561) and the NIH/NIDDK-funded Boston-Area DERC (P30DK057521) for core services. Some strains were provided by the CGC, funded by the NIH Office of Research Infrastructure Programs (P40OD010440), and the *C. elegans* Knockout Consortium. Figures 1A, 3C, and 7A were created with BioRender.com.

## Supplementary material

### Supplementary file descriptions

**Supplementary file 1 (separate file)**. Tabular and survival data including 3 biological replicates (unless otherwise noted) are shown for lifespan experiments related to Figures 1, 3, 4, 5, Figure 1-figure supplement 1, Figure 4-figure supplement 1, and Figure 5-figure supplement 1. Data present a summary of the conditions tested which, if applicable, include: (1) drug treatment with vehicle control and 4.5 mM phenformin or 50 mM metformin and/or (2) RNAi treatment to knockdown expression of the specific denoted gene. The *C. elegans* strain, number of subjects, restricted mean (days), standard error, 95% confidence interval (C.I.), 95% median C.I., and *P*-values for relevant comparisons are noted amongst all conditions. ns, not significant; *, *P* < 0.05; **, *P* < 0.01; ***, P *<* 0.001; ****, *P* < 0.0001 by log-rank analysis.

**Supplementary file 2 (separate file)**. Excel file containing raw, normalized, and normalized and log_10_ transformed mass spectrometry data for phosphatidylethanolamine containing ether lipids detected by LC-MS/MS. Data from **3** biological replicates are shown for molecules indicated for vehicle or 4.5 mM phenformin treatment, for 4 different genetic backgrounds: wild type animals (N2), BX10 (*ads-1* mutant), BX259 (*acl-7* mutant) and BX275 (*fard-1* mutant). Compound identity for each detected lipid as well as raw, normalized or transformed mass counts on each of three tabs. Note, several of the lipids were not uniformly detected or of low abundance, and thus were filtered by the MetaboAnalyst parameters used and not represented on the “Normalized” and “Normalized-Log10 Transformed” tabs.

**Figure 1—figure supplement 1.**
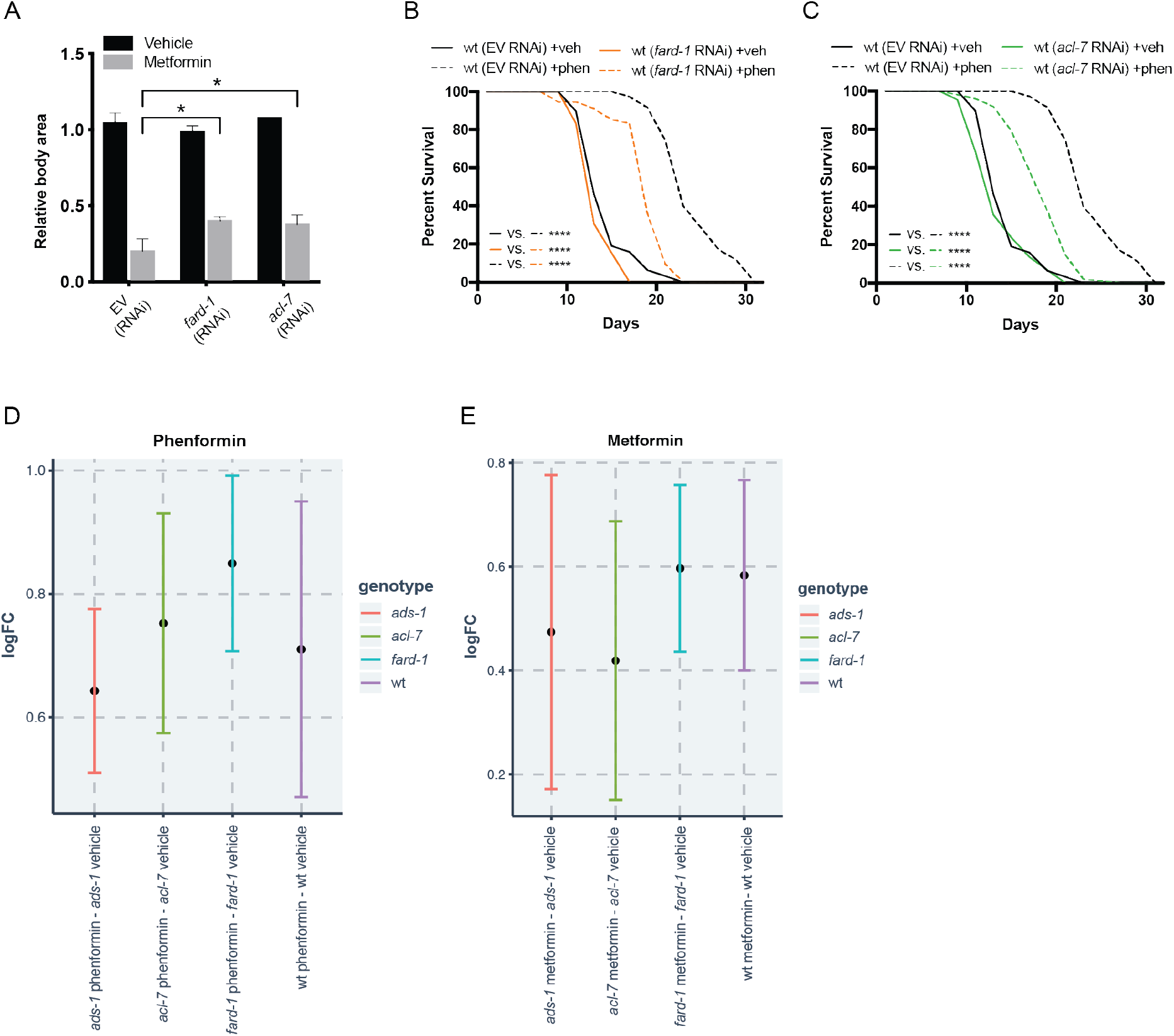
Reduced function of genes responsible for ether lipid biosynthesis partially suppresses biguanide effects of growth and lifespan without affecting biguanide levels. (A) RNAi to *fard-1* and *acl-7* induce *C. elegans* resistance to growth inhibition by 160 mM metformin treatment. *, *P* < 0.05, by two-way ANOVA, n = 2 biological replicates. (B-C) RNAi knockdown of *fard-1* (*B*) and *acl-7* (*C*) in *C. elegans* partially suppresses phenformin’s effect on lifespan extension. For *B* and *C*, results are representative of 3 biological replicates. ****, *P* < 0.0001 by log-rank analysis; for tabular survival data and biological replicates see also Supplementary file 1. (D) Log fold change (LogFC) of phenformin abundance in samples treated with 4.5 mM phenformin versus vehicle reveals that the increase in phenformin levels in wild type and three ether lipid deficient mutants is similar. (E) LogFC of metformin abundance in samples treated with 50 mM metformin versus vehicle show that metformin increases are similar across all 4 strains (*E*). Bars represent mean and 95% confidence intervals.

**Figure 2—figure supplement 1.**
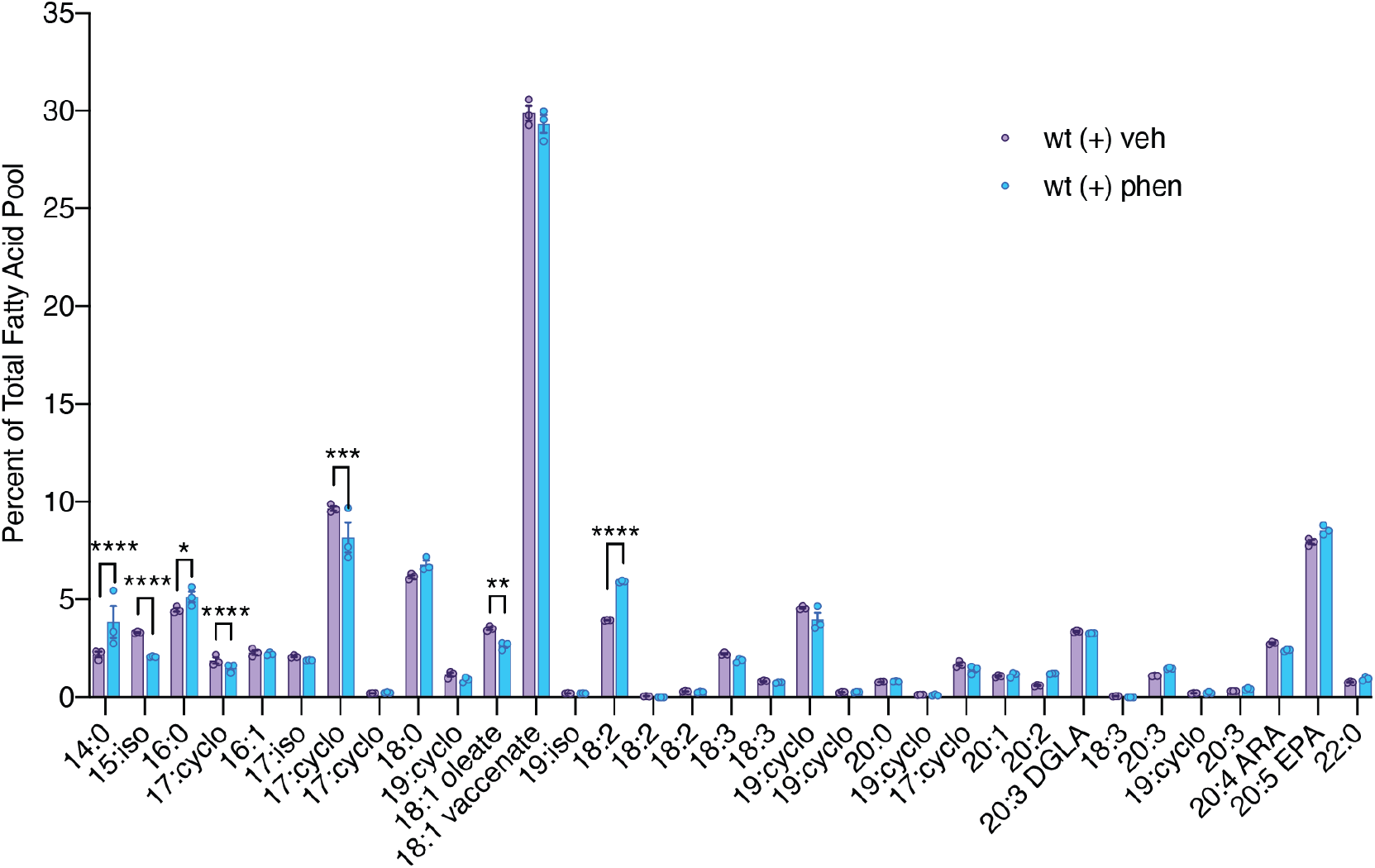
Biguanide treatment modulates abundance of fatty acids in *C. elegans*. A comparison of the percent of the total fatty acid pool for 33 fatty acids shows that 7 fatty acids are significantly altered in phenformin treated wild type worms. n = 3 biological replicates. *, *P* < 0.05; **, *P* < 0.01; ***, *P* < 0.001; ****, *P* < 0.0001 by multiple t-tests (corrected for multiple hypothesis testing with two-stage step-up method of Benjamini, Krieger, and Yekutieli).

**Figure 2—figure supplement 2.**
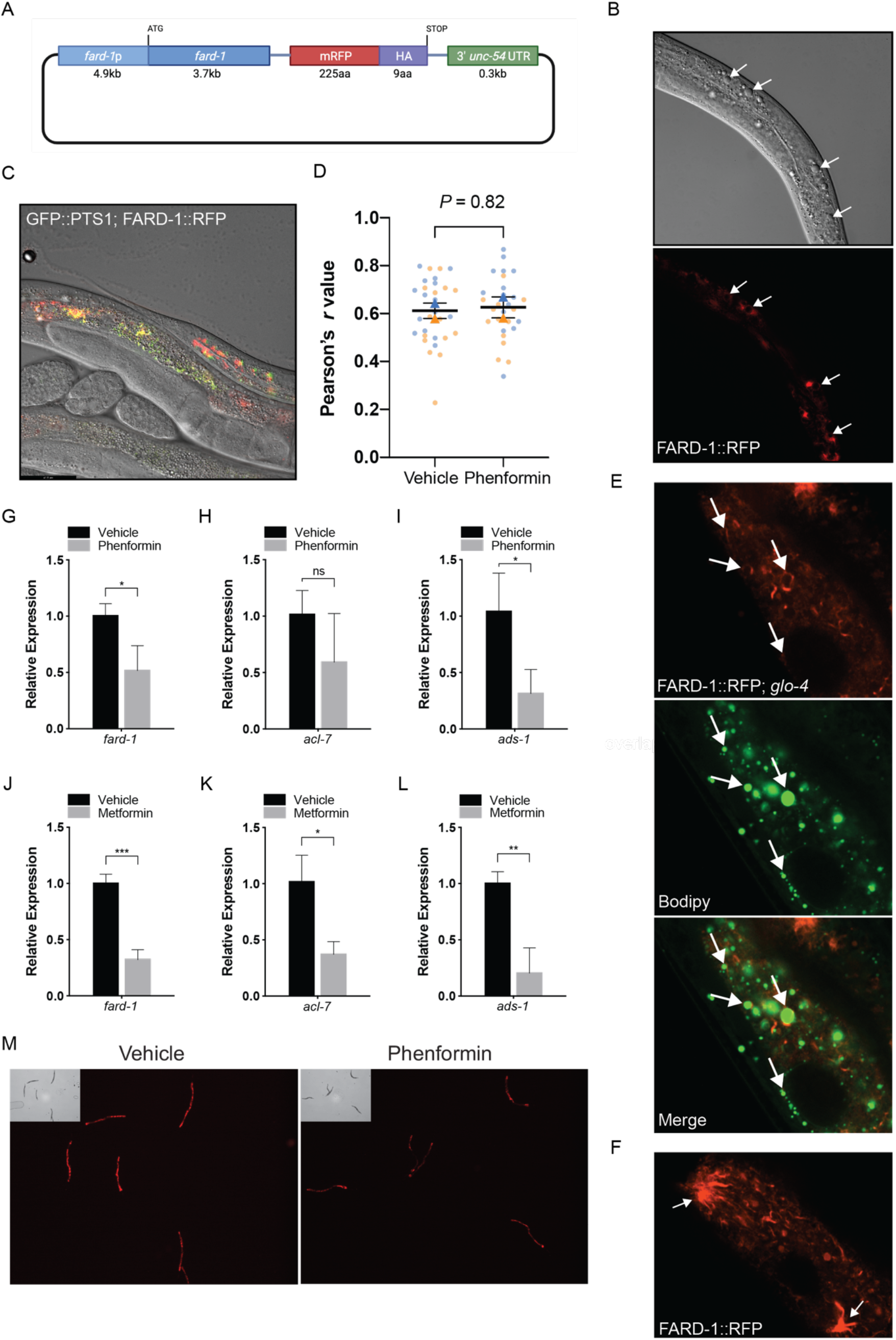
FARD-1::RFP localizes to intestinal lipid droplets and peroxisomes and is not positively regulated at the RNA or protein level by phenformin. (A) Diagram of the *C. elegans* FARD-1::RFP overexpression reporter. (B) FARD-1::RFP *(fard-1 oe1)* exhibits intestinal expression in *C. elegans*. FARD-1 displays a cytoplasmic distribution and an association with structures resembling lipid droplets (B, *arrows*). (C) Co-expression of FARD-1::RFP and peroxisomally targeted GFP::PTS1 in transgenic animals indicates partial colocalization of FARD-1 with peroxisomes in intestine. (D) Superplot displays colocalization of RFP and GFP in vehicle or phenformin treated GFP::PTS1; FARD-1::RFP transgenics (N= 20 total worms assessed; 5 worms per condition; 3 images per worm (upper/mid/lower intestine)) for a total of 15 images (dots) per replicate; blue = replicate 1, orange = replicate 2). Correlation coefficients were separately calculated for each biological replicate and the mean is represented for each pool (blue or orange triangle). These two means were then used to calculate the average (horizontal bar), standard error of the mean (error bars), and P value. Analysis of the average Pearson’s r values demonstrates no significant difference between colocalization of FARD-1::RFP and GFP::PTS1 in vehicle or phenformin-treated worms. n = 2 biological replicates. (E) Confocal imaging of an integrated FARD-1::RFP reporter *(fard-1 oe3)* in *C. elegans* stained with C1-BODIPY-C12 (treated with *glo-4* RNAi to remove BODIPY positive lysosome related organelles) demonstrates localization of FARD-1 protein to the surface of lipid droplets in the worm intestine. (F) In *fard-1(oe3)* transgenics, confocal imaging indicates FARD-1::RFP organization into web-like structures and bright punctae that represent the intersection of these “webs”. These structures may represent smooth endoplasmic reticulum. (G-I) Levels of *fard-1, acl-7*, and *ads-1* mRNA decrease in wild type *C. elegans* treated with 4.5 mM phenformin versus vehicle. n = 3 biological replicates; ns, not significant; *, *P* < 0.05 by unpaired *t*-test. (J-L) Levels of *fard-1, acl-7*, and *ads-1* mRNA decrease in wild type *C. elegans* treated with 50 mM metformin versus vehicle. N = 3 biological replicates; *, *P* < 0.05; **, *P* < 0.01; ***, *P* < 0.001 by unpaired *t*-test. (M) Phenformin (4.5 mM) results in decreased expression of the FARD-1::RFP translational reporter (*fard-1 oe1*). n = 3 biological replicates; total assessed: N = 30 worms per condition (10 worms per replicate).

**Figure 4—figure supplement 1.**
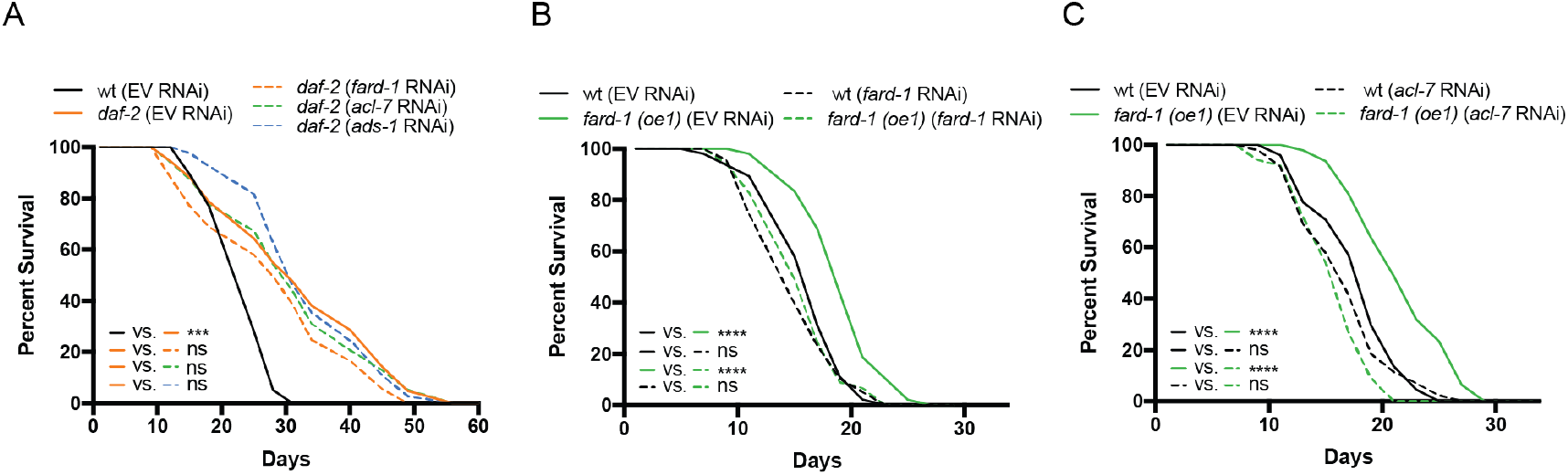
Ether lipid biosynthetic genes are not necessary for *daf-2*-dependent lifespan extension, and *fard-1* overexpression extends lifespan in a manner dependent upon ether lipid biosynthesis. (A) *daf-2* mutants display extended lifespan relative to wild type animals. RNAi knockdown of *fard-1, acl-7*, and *ads-1* does not impact lifespan extension in these mutants. (B-C) RNAi knockdown of *fard-1* (*B*) and *acl-7* (*C*) suppresses *fard-1* overexpression*(oe1)*-associated lifespan extension. For A-C, results are representative of 2-3 biological replicates. ns, not significant; ***, *P* < 0.001; ****; *P* < 0.0001 by log-rank analysis. For tabular survival data and biological replicates see also Supplementary file 1.

**Figure 5—figure supplement 1.**
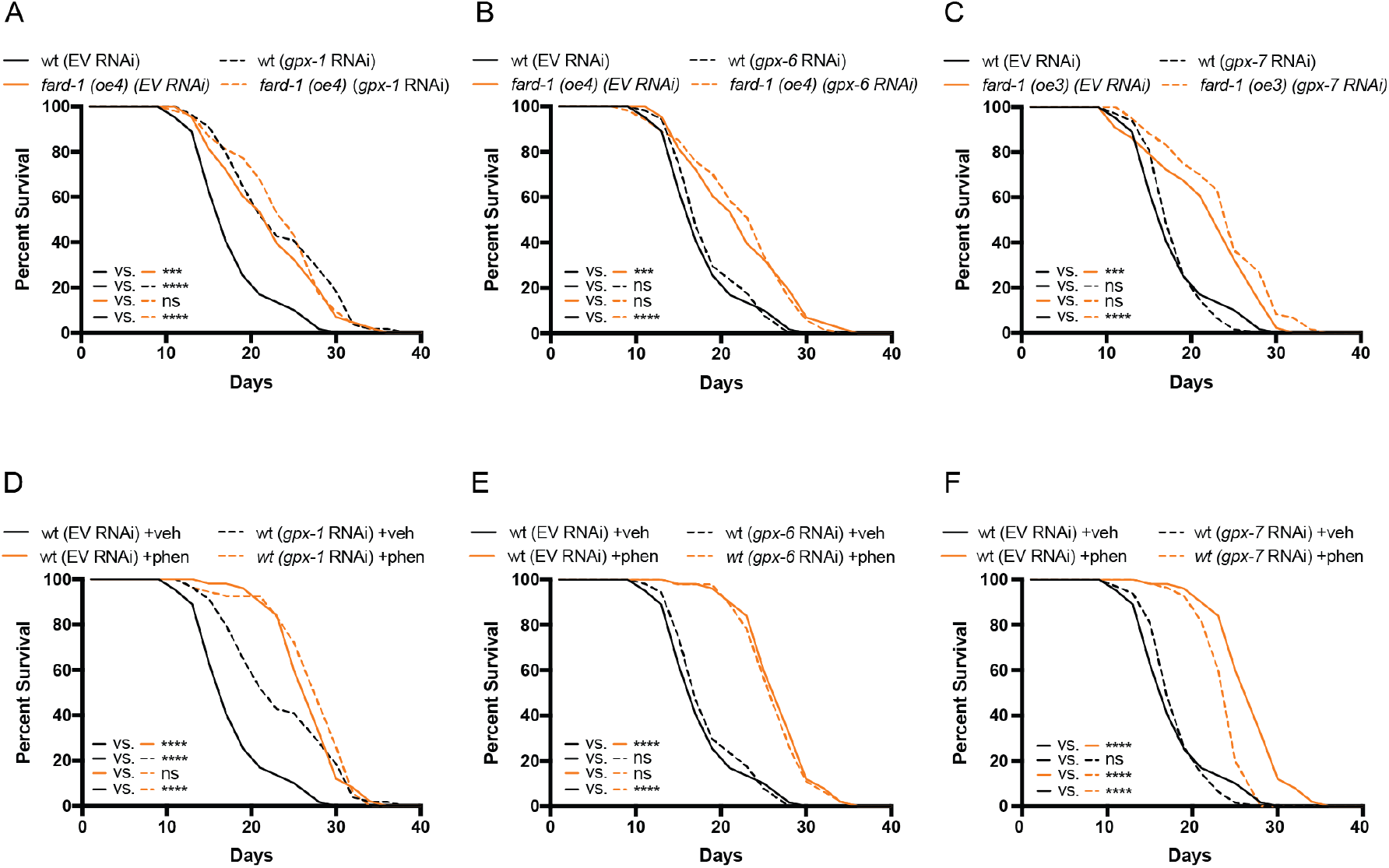
Genetic induction of ferroptosis does not impact *fard-1* overexpression nor biguanide-mediated lifespan extension. (A-C) Independent knockdown of glutathione peroxidases, *gpx-1* (A), *gpx-6* (B), or *gpx-7* (C) by RNAi does not mitigate lifespan extension by integrated *fard-1* overexpression *(fard-1 oe3* and *fard-1 oe4)*, as would be expected if fard-1 overexpression extended lifespan by lowering ferroptosis. *gpx-1* unexpectedly extends lifespan in a non-additive manner with *fard-1(oe)*. (D-F) Similarly, knockdown of *gpx-1* (D), *gpx-6* (E), or *gpx-7* (F) by RNAi do not suppress phenformin-mediated lifespan extension. For *A-F*, results are representative of 2-3 biological replicates. ***, *P* < 0.001; ****, *P* < 0.0001 by log-rank analysis. For tabular survival data and biological replicates see also Supplementary file 1.

